# Perinatal Elimination of Genetically Aberrant Neurons from Human Cerebral Cortex

**DOI:** 10.1101/2024.10.08.617159

**Authors:** Diane D. Shao, Yifan Zhao, Joseph Brew, Yahia Adla, Urmi Ghosh, Vinayak V. Viswanadham, Lovelace J. Luquette, Sijing Zhao, Andrea J. Kriz, Xuyu Qian, Takumi Taketomi, Shannon Coy, Sandro Santagata, Michael Miller, Fuminori Tsuruta, Peter J. Park, Christopher A. Walsh

**Affiliations:** Department of Neurology, Boston Children’s Hospital, Boston, MA, USA; Department of Pediatrics, Division of Genetics and Genomics, Boston Children’s Hospital, Boston, MA, USA; Department of Biomedical Informatics, Harvard Medical School, Boston, MA, USA; Program in Health Sciences & Technology, Harvard Medical School & Massachusetts Institute of Technology, Boston, MA, USA; Current address: Wellcome Sanger Institute, Hinxton, UK; European Bioinformatics Institute, European Molecular Biology Laboratory, Hinxton, UK; Howard Hughes Medical Institute, Chevy Chase, MD, USA; Master of Public Health Program, School of Public Health, Brown University, Providence, RI, USA; Biological and Biomedical Sciences Graduate Program, Harvard Medical School, Boston, MA, USA; Department of Pediatrics, Children’s Hospital of Philadelphia; Perelman School of Medicine, University of Pennsylvania, Philadelphia, PA, USA; Ph.D. Program in Human Biology, University of Tsukuba, Tsukuba, Ibaraki, Japan; Department of Pathology, Brigham and Women’s Hospital, Boston, MA, USA; Laboratory of Systems Pharmacology, Harvard Medical School, Boston, MA, USA; Ludwig Center, Harvard University, Boston, MA, USA; Institute of Life and Environmental Sciences, University of Tsukuba, Tsukuba, Ibaraki, Japan

## Abstract

Human neurons are postmitotic and long-lived, requiring precise genomic regulation to maintain function over a lifetime. Normal neuronal function is highly dependent on gene dosage, with copy number variants (CNVs) and heterozygous point mutations associated with a host of neurodevelopmental and neuropsychiatric conditions [1–5]. Here, we investigated the landscape of somatic CNVs arising in fetal human brains, and how they change over development, to understand the processes that generate normal neuronal genomes. We sequenced 2,158 single neurons from human postmortem brain using two distinct single cell whole genome sequencing (scWGS) approaches. Tn5-transposase based (TbA) scWGS of 1,327 neurons from 16 individuals ranging in age from gestational week 14 to 90 years old resulted in 8,765 CNVs. Primary template amplification (PTA) was used to assess for CNV from 831 neurons from 12 individuals. Up to 46% of neurons in prenatal cortex showed aberrant genomes, characterized by widespread CNVs of multiple chromosomes, but this proportion reduces by 4-5 fold after birth across the two scWGS approaches (p=7.5×10^−6^, Fisher meta-analysis). We identified micronuclei in the developing cortex *in situ*, reflecting chromosomal material missegregated during neurodevelopment [6–8]. Neurons with widespread CNVs were eliminated in the perinatal period, while neurons with smaller CNV burden slowly declined during postnatal aging. CNVs in surviving neurons were depleted for genes that are dosage-sensitive or involved in neurodevelopmental disorders (p*<*0.05), suggesting selective elimination of neurons with CNVs involving these critical genes. We surveyed 44,861 nuclei with 10X Genomics scATAC/RNAseq and determined that neurons with high CNV burdens also showed abnormal expression of synaptic gene sets, suggesting that abnormal synaptic gene regulation contributes to neuronal elimination. Elimination of defective neuronal genomes during synaptogenesis may represent a critical process of genome quality control and a vulnerable target of factors that contribute to neurodevelopmental disease.

## Introduction

Genome-scale copy number variants (CNV) are common in early post-zygotic development – with up to 73% of embryos exhibiting large chromosomal aneuploidy [7, 9–11]. Similarly, human neural progenitor cells (NPCs) display extensive mosaic aneuploidy, involving up to 30% of progenitors [12, 13]. Rates of aneuploidy in adult neurons have been controversial, with some reports of widespread aneuploidy [14], while other reports suggest rare aneuploidy [15–19], or frequent, predominantly sub-chromosomal CNVs [17, 20, 21]. These discrepancies arise from differences in methodology and the sensitivity of techniques, making it unclear whether the high aneuploidy rates observed early in development are maintained through to adulthood in the brain.

The control of copy number is especially crucial in brain, since genetic variation in neurons can have profound functional impact. Autism, schizophrenia, epilepsy, intellectual disability and other neuropsychiatric conditions are all commonly caused by chromosomal or sub-chromosomal CNV, as well as by heterozygous loss of function point mutations in more than one thousand genes essential for brain functions [2, 3, 22, 23]. Many of these genes encode structural proteins of the synapse, or regulators of synaptic plasticity, suggesting that key aspects of neuronal plasticity underlying these conditions are intensely sensitive to gene dosage [24–26].

While a variety of cell cycle checkpoints regulate progenitors to prevent or eliminate cells with aberrant genomes [14, 27–29], it is not clear whether any such pathways might exist to control gene dosage in noncycling cells like neurons. One hypothesis suggests that genetically aberrant neurons are eliminated through “programmed cell death” [28, 30], a form of widespread apoptosis which in human cerebral cortex is estimated to remove 20-50% of neurons during development [27, 31]. In all examined mammalian species, the most dramatic wave of apoptosis eliminates *∼*30% of neurons in the perinatal period [32], followed by ongoing elimination of postnatal neurons [28, 33] coincident with the establishment of neuronal circuits and connectivity mediated by synapse formation and pruning [34]. While roles for synapse formation in the regulation of neuronal survival have been described for more than a century [35, 36], the mechanisms and significance of widespread neuronal loss have not been clear, since dying neurons tend to be dispersed across the cortex, without clear relationship to neuronal cell type, laminar location, or topographic patterns of connectivity [28, 33, 37].

Using single-cell whole genome sequencing (scWGS), we compared copy number variants (CNVs) in postmitotic neurons during neurodevelopment and aging. Our analysis shows widespread CNVs across chromosomes in fetal neurons, followed by removal of most neurons with highly defective genomes around the time of birth. We also confirmed prior findings of a slower removal of defective cells occurring postnatally into aging [20]. Our data also show that this genome quality control mechanism is linked to gene pathways essential for synapse development, including disease-relevant genes. These findings suggest that synaptogenesis plays a key role in the control of genome quality in neurons, acting most dramatically near birth but also extending into later life. This process is potentially vulnerable to environmental or genetic defects that disrupt normal brain development.

## Results

### Accurate copy number detection in single neurons identifies aberrant genomes in cortical neurons

To profile CNVs in the human prenatal brain, we first generated low-coverage scWGS data from 1,327 neurons, collected from non-diseased postmortem brains from (1) nine prenatal donors ranging from gestational weeks 14-25 and (2) seven postnatal brains from infancy to 90 years (Fig. 1a, Table S1). To obtain a pure population of postmitotic prenatal cerebral cortical neurons, we cryo-dissected the human prenatal cortical plate (Fig. 1b) followed by flow cytometry to isolate single neuronal nuclei. Dissection of cortical plate neurons provides an exclusively postmitotic, noncycling fetal neuronal population, because NPCs and glial nuclei are located in undissected deeper cortical regions. Neurons migrate into the cortical plate after withdrawing from the cell cycle and becoming permanently postmitotic. Single-nucleus RNA sequencing (snRNA-seq) confirmed that our cortical plate dissection yielded 98% purity for neurons (Fig. 1b-c). Postnatal nuclei suspensions were stained with anti-NeuN antibody (neuronal marker) and DAPI prior to flow cytometry to isolate neuronal nuclei as previously described [38]. scWGS libraries were generated using Tn5-based amplification (TbA) after removal of chromatin-associated proteins with Proteinase K and sequenced at *∼*0.05X depth (Extended Data Fig. 1a; Methods). Tn5-based single-cell whole genome amplification (scWGA) [39–41] for scWGS results in even coverage of the genome, i.e., low Median Absolute Pairwise Difference (MAPD) scores, and is cost-effective for scaling. In our case, this scWGA method was chosen to enable a bioinformatic validation pipeline to overcome typical challenges in validation of CNV, as the original single cell DNA substrate is destroyed in the process of scWGA. To establish a reference for high-quality genomic profiles, we characterized cultured neuronal progenitor cells (NPCs) with and without disrupted nuclear integrity after inducing cell-death with staurosporin (Extended Data Fig. 1b), and applied these data as the basis for transfer learning in order to distinguish high-quality from poor-quality sequencing profiles from our postmortem tissues (Extended Data Fig. 1c-h; Methods). These stringent quality control steps together with restricting MAPD to less than 0.5 (Extended Data Fig. 1i) resulted in 948 high-quality single-cell neuronal genomes.

**Fig. 1:**
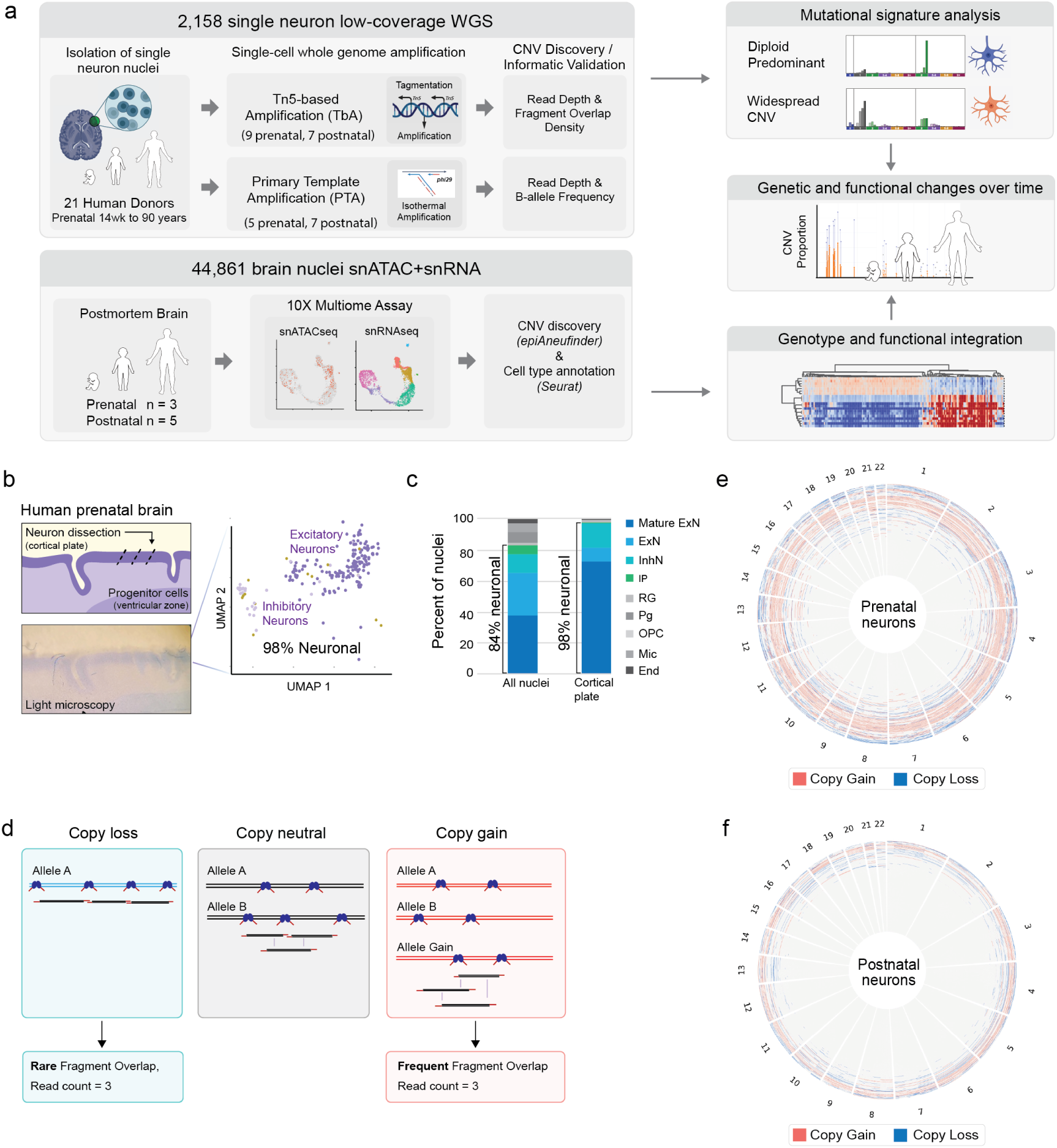
A subpopulation of human postmitotic neurons harbors aberrant copy number patterns. **a.** Overview of study design for scWGS, copy number calling, and downstream analyses. **b.** Cortical plate dissection of frozen normal prenatal human brain isolates neuronal nuclei, confirmed by single-nucleus RNA sequencing. Neuronal nuclei (dark and light purple); non-neuronal nuclei (yellow). **c.** Nuclei isolation and confirmation of cell types from prenatal cortical plate dissection. Excitatory neuron (ExN), inhibitory neuron (InhN), intermediate progenitor (IP), radial glia (RG), neural progenitor (Pg), oligodendrocyte progenitor cell (OPC), microglia (Mic), endothelial cell (End). **d.** Schematic illustration of Tn5-based amplification libraries in which single-genomes create distinct patterns of molecular overlap that distinguishes copy number states. **e.** Circos plot of CNV profiles and gain/loss distribution in 470 high-quality prenatal human cortical neurons. Chromosomes are arranged circumferentially. Neurons are ordered based on total CNV sizes. **f.** Circos plot of CNV profiles and gain/loss distribution in 478 high quality postnatal human cortical neurons. Chromosomes are arranged circumferentially. Neurons are ordered based on total CNV sizes.

To identify CNVs, we utilized HiScanner [42], an algorithm we previously developed specifically for CNV detection in scWGS data. For robustness against amplification noise and sparse sequencing coverage, HiScanner models technology-specific allelic dropout patterns from scWGS data and leverages probabilistic segmentation to maintain high detection precision [43, 44]. In addition, we designed another informatic validation approach: Fragment Overlap Density (FOD) metric. Tn5-transposase-based libraries result in a specific library property in that we expect all amplified fragments to retain the start and end position of the original tagmented molecule; thus, in a copy-loss region where only one allele is present, no fragments should overlap, unlike in copy-neutral or copy-gain regions (Fig. 1d). This heuristic is represented by the FOD metric (Methods), which we use to further increase our CNV detection accuracy (Supplemental Results; Extended Data Fig. 2a-g).

Surprisingly, we found that neurons with aberrant genomes were present in both the prenatal and postnatal human cerebral cortex (Fig. 1e-f, Table S2). Many genomes with alterations had multiple affected regions across more than one chromosome, often involving large copy number changes. In prenatal samples, copy losses were nominally more frequent than copy gains, with 53% (3090/5844) of CNVs being copy losses, and 47% (2754/5844) copy gains. In postnatal neurons, copy losses (59%, 1722/2921) were again more frequent than copy gains (41%, 1199/2921).

We observed that the mean percentage of neurons with at least 1 CNV of any size is significantly higher in prenatal compared to postnatal neurons, 62.9% and 29.2%, respectively (p-value =0.002; Student’s t-test, two-tailed; Extended Data Fig. 2h). This perinatal difference was significant for the percentage of neurons with copy gains (40.6% prenatal to 10.0% postnatal, p-value = 0.0002, Student’s t-test, two-tailed) as well as copy losses (43.9% prenatal; 23.4% postnatal, p-value = 0.04, Student’s t-test, two-tailed).

### Neurons with widespread CNVs decrease between mid-gestation and postnatal life

Because arbitrary thresholds based on CNV size or count fail to capture the true biological complexity [45] of each neuron, which may include both small and chromosome-scale CNVs, we applied mutational signature analysis [46, 47] to classify cells with aberrant genomes. Based on the COSMIC copy number signature catalog [47], we classified neurons into four distinct “signature groups”: One group – *Diploid Predominant CNV neurons* (COSMIC SigCN1 signature) – represents neurons with primarily diploid genomes and those with minimal CNVs (Fig. 2a). Three groups represent neurons with “Widespread CNV”: *Gain Predominant* (combination of SigCN1 and SigCN2), *Focal Loss Predominant* (SigCN1 and SigCN9), and *Broad Loss Predominant* (SigCN1 and SigCN13) (Fig. 2a, Extended Data Fig. 3). Widespread CNV neurons, compared to Diploid Predominant CNV neurons, harbor genomic alterations that encompass larger regions of the genome, larger size distribution of CNV, and have more aneuploidy (here, defined as gain or loss of *>*90% of a chromosome) (Extended Data Fig. 4). The neuronal copy number signatures showed high resemblance to those defined in COSMIC from cancers (mean cosine similarity 0.9716; Extended Data Fig. 3d), suggesting that neuronal CNVs may have arisen in dividing cells, presumably NPCs, in the developing human brain.

**Fig. 2:**
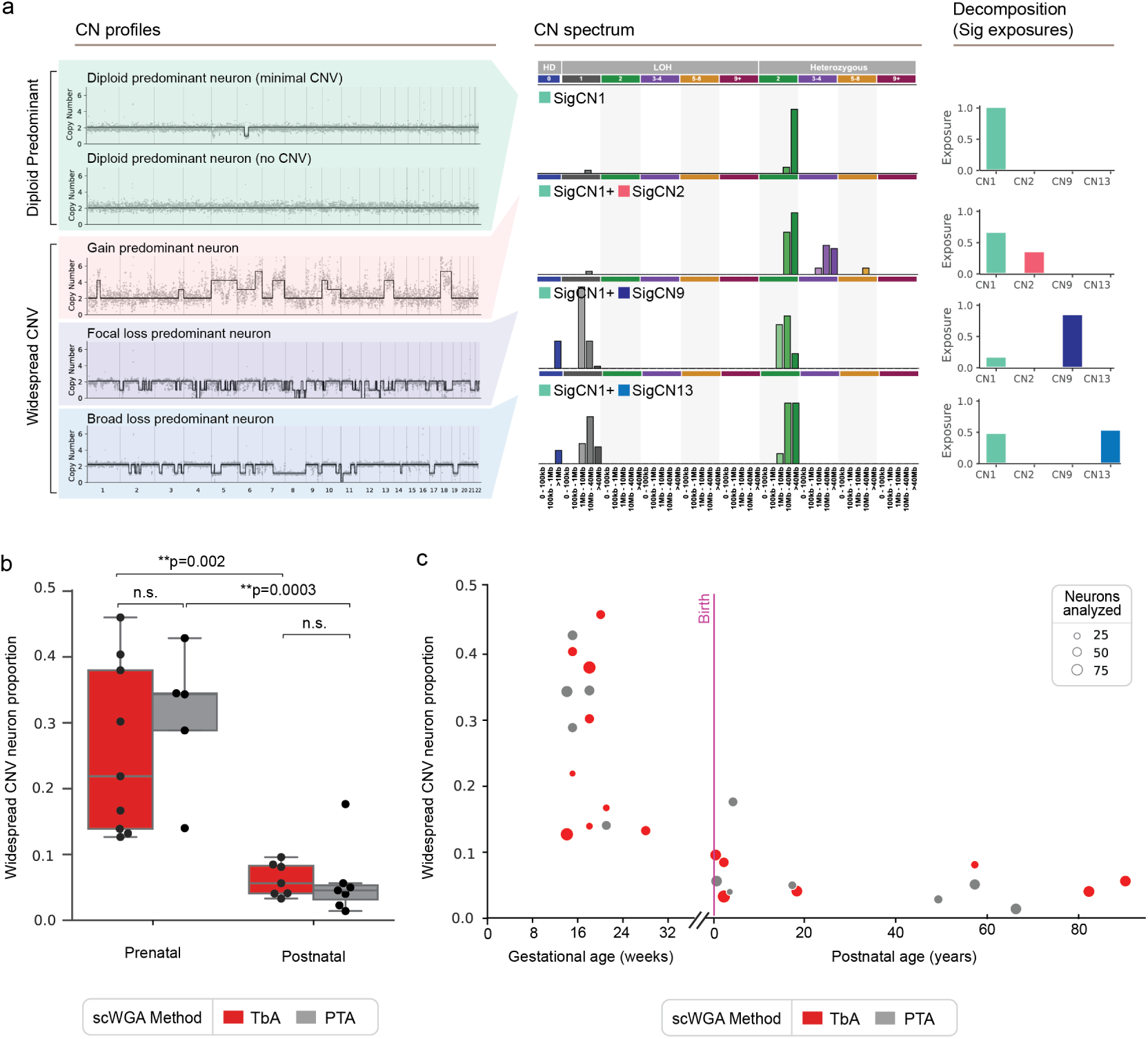
Widespread CNVs that predominate prenatally are associated with copy number signatures. **a.** Copy number signatures represented by the four primary signature clusters identified across all human neurons. The left panel shows representative single-neuron copy number profiles for each of the signature groups, the middle panel shows the composite copy number spectrum characteristic of each group, and the bar plots in the right panel shows the signature exposure decomposition. **b.** Proportion of neurons with widespread CNVs observed in human prenatal versus postnatal neurons determined using two distinct scWGA technologies. Box plots indicate the median, first and third quartiles (hinges) and the most extreme data points no farther than 1.5X IQR from the hinge (whiskers). IQR, interquartile range. p, p-value, two-tailed Student’s test. **c.** Frequency of neurons with widespread CNVs rapidly decreases between midgestation and the postnatal period, and remains low throughout postnatal life. Each point represents neurons analyzed for a single donor. Results from two distinct scWGA technologies are shown.

The proportion of neurons with widespread CNVs (last 3 of the 4 groups defined above) in prenatal samples was on average 25.9% (range 12.7-46.0%), but decreased significantly to *∼*6.2% postnatally (p=0.002; Student’s t-test, two tailed) (Fig. 2b), 4.2 fold. The mean during infancy (0-2 years) was 7.1% and was 4.8% for elderly samples (*>*60 years) (Fig. 2b-c), suggesting relative stability of this population during postnatal aging.

To validate our results from TbA, we performed primary template amplification (PTA) [48] in single neurons, as an orthogonal scWGA method, on an additional 831 neuronal nuclei from 5 prenatal and 7 postnatal donors, yielding 659 high quality neurons that passed QC filters (Table S1-S2; Extended Data Fig. 5). Neurons amplified with PTA were sequenced at higher depth (*∼*0.5X), resulting in generally even genome amplification as determined by MAPD (Extended Data Fig. 5a-b). Copy number signatures derived from PTA-amplified neurons also segregated into distinct groups highly concordant with the Diploid Predominant and Widespread CNV categorizations from TbA (Extended Data Fig. 5c-f). The mean widespread CNV neuron proportion decreased 5-fold between the prenatal to postnatal periods, from 30.1% to 5.8% of neurons, respectively (p = 0.0003, Student t-test, two-tailed; Fig. 2b-c), and for each age category was not significantly different from the widespread CNV proportion from TbA. The combined Fisher meta-analysis *p*-value across both scWGA methods for the prenatal-postnatal changes was *p* = 7.5 *×* 10*^−^*^6^. To ensure that the observed reduction in the proportions of neurons with aberrant genomes are not affected by poor quality cells, we checked that there is no significant association of widespread CNV proportion with the percentage of neurons passing QC filters and sample size (Extended Data Fig. 6a-b), nor with postmortem interval (Extended Data Fig. 6c).

Prior reports indicate that the proportion of neurons harboring any CNV declines with age [17, 20]. We thus also analyzed our data, including both TbA- and PTA-amplified neurons, based on the proportion of neurons harboring at least 1 CNV. We noted again that the primary decrease in CNV proportion from 63.9% to 30.9% occurs between the prenatal and early postnatal periods (age 0-5 years)(p=0.0007, Student t-test with Bonferroni correction; Extended Data Fig. 7a). From early postnatal to later ages (6+ years), we found a nominal trend towards decrease in mean proportion from 30.9% to 22.5% (p=0.27, Student t-test, Bonferroni correction), indicating an underpowered analysis from our data alone. Therefore, we performed a meta-analysis, combining data from the two scWGA technologies in our study and 7 additional scWGA studies in postnatal neurons using a linear mixed-regression model, thus accounting statistically for differences in sequencing platform and CNV calling methods from independent studies (Table S3). At birth, the model estimated *∼*29.4% of cells carry a CNV (intercept p*<*0.0001), and age was significantly negatively associated with the proportion of neurons carrying any CNV (p*<*0.0001)(Extended Data Fig. 7b), supporting an ongoing elimination of CNV-carrying neurons during postnatal life.

### Neuronal CNVs result from genomic stress in progenitor cells

To determine whether CNVs arise in replicating progenitor cells or postmitotic neurons, we analyzed their associations with DNA replication timing [49, 50], and physiological double strand breaks (DSBs) [51, 52]. We found that CNV breakpoint density was highly correlated with replication timing (Pearson r=-0.95, p*<*0.001), such that CNV breakpoints were more frequent in early-replicating regions (Fig. 3a-b). This result suggests that CNVs may be generated in mitotic progenitor cells during DNA replication.

**Fig. 3:**
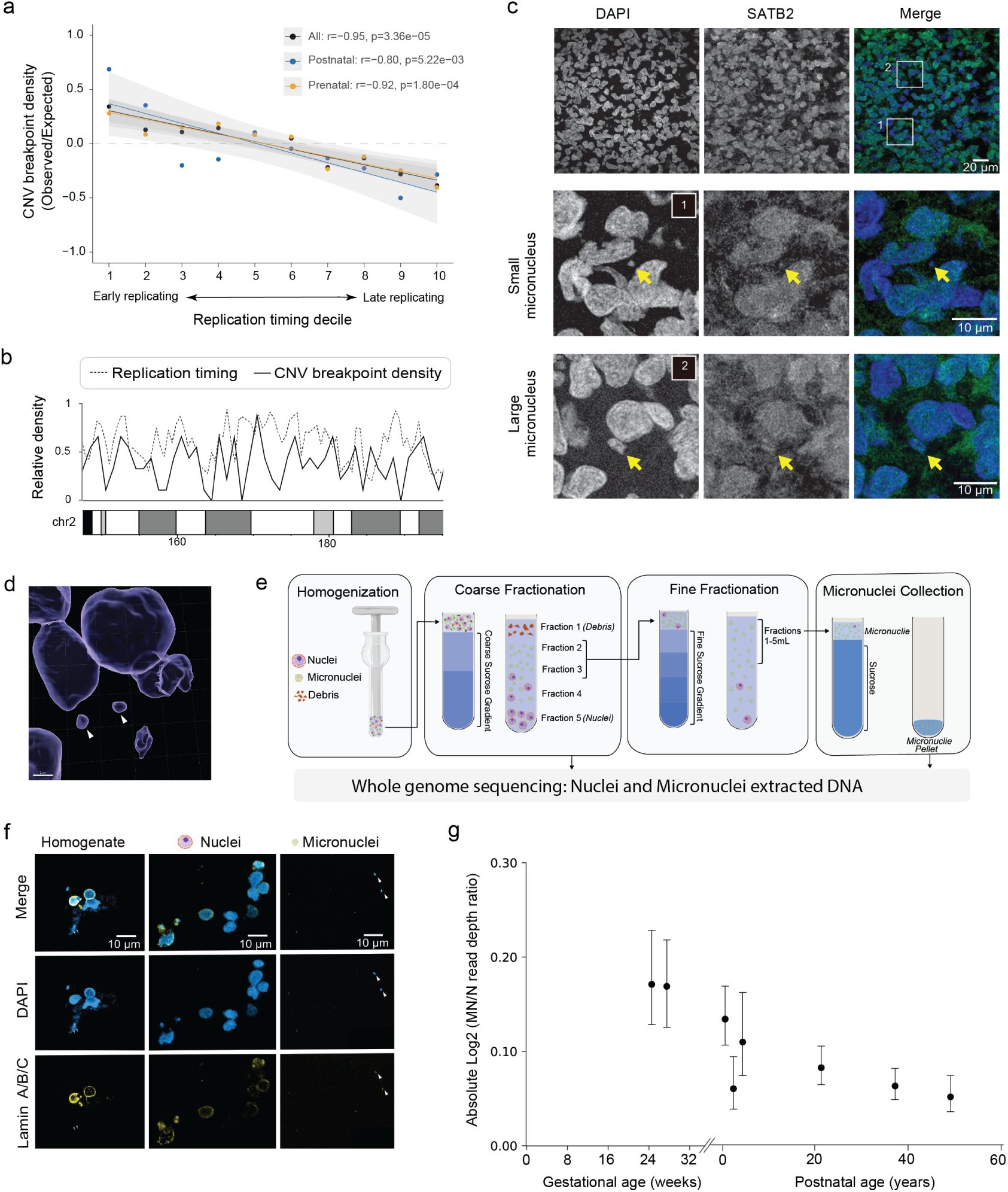
Etiology of CNV breakpoints. **a.** Association between CNV breakpoints and replication timing deciles. **b.** Representative region on chromosome 2 showing local association between density of CNV breakpoints and replication timing. **c.** Micronuclei in the cortical plate are associated with neuronal nuclei, observed by confocal microscopy and marked by SATB2 staining (prenatal neurons) and DAPI staining (DNA). **d.** 3D reconstruction of DAPI staining from confocal microscopy of the cortical plate showing primary nuclei and micronuclei (arrowheads). **e.** Schematic of serial sucrose gradient fractionation to obtain micronuclear and primary nuclear fractions for sequencing. **f.** Immunohistochemistry of homogenate, nuclei fraction, and micronuclei obtained from serial sucrose gradient fractionation to visualize content of each fraction. Nuclei and micronuclei (arrowhead) stain positive for both nuclear lamin and DAPI. **g.** Absolute value of micronuclear fraction of genomic content relative to paired nuclear fractions shows reduction with age. Each point represents the average of autosomes for each donor evaluated, and lines indicate 95% confidence intervals. Data were adjusted for chromosomal GC content by linear regression.

We next evaluated hotspots of DSBs in NPCs determined previously by in-suspension Breaks Labeling In Situ and Sequencing (BLISS) [51], as DSBs have been associated with CNV generation in neurological disease [52]. Interestingly, BLISS hotspots (top 1% of BLISS regions) were associated with copy loss at recurrent CNV breakpoints between donors and recurrent CNV breakpoints associated with copy gains showed a non-significant trend toward enrichment at BLISS hotspots (p=0.06; Extended Data Fig. 8a). In contrast, CNV breakpoints unique to individual neurons (gain or loss) were not significantly associated with BLISS hotspots (Extended Data Fig. 8b). Analysis of within-donor recurrent breakpoints was insufficiently powered. Together, these findings suggest that DSB-prone loci contribute to recurrent CNV breakpoints, with the strongest evidence for copy losses. DNA replication stress has been associated with the formation of micronuclei, which encapsulate missegregated chromosomes or fragments that lie outside of the nucleus, as observed in both early postzygotic cell divisions and cancer [7, 53, 54]. Thus, we hypothesized that micronuclei would be visible in some human prenatal neurons, as they have also recently been observed in mice [55]. High-resolution microscopy on fixed slices of human fetal brain at mid-gestation, stained with DAPI, revealed co-occurrence of micronuclei in cells with neuronal identity as assayed by immunostaining with the prenatal neuronal marker, SATB2 (Fig. 3c). 3-D reconstructed images demonstrate localization of micronuclei in the fetal cortical plate (Fig. 3d). We observed that micronuclei were associated with *∼*6% of primary nuclei in the cortical plate (Table S4), providing direct, *in situ* demonstration of relatively common, missegregated chromosomal material at midgestation.

We sought to orthogonally confirm the elimination of cells with widespread CNVs over time by evaluating whether micronuclei are also cleared over time (Fig. 3e-g). We isolated micronuclear and nuclear fractions from human brain tissue from fetal, infant, and elderly individuals by serial gradient sucrose fractionation [56] with visualization to identify micronuclei-enriched genetic content (Fig. 3e-f). By comparing the genetic content of micronuclear and nuclear fractions using genome sequencing, we found that micronuclei in the prenatal samples harbored aberrant genetic material, observed as elevated read depth relative to the nuclear fraction (Extended Data Fig. 9, Table S5). The magnitude of this deviation decreased in infancy/childhood and old age, indicating clearance of micronuclei genetic material over time (Extended Data Fig. 9a). The significant decrease in micronuclear content with age persisted after regression analysis adjusting for chromosome size and GC content (Fig. 3g; Supplemental Methods). The age-related decline in micronuclear content occurs temporally in parallel with the decline in CNVs in primary nuclei (as measured by TbA and PTA), providing further evidence for the selective depletion of genomically aberrant neurons over time.

### Neuronal CNVs disrupt gene modules relevant to synaptic signaling and maintenance

In order to identify pathways regulating the elimination process by associating CNV changes in single neurons with gene expression, we performed 10X multiomic snATAC/RNA-seq of 44,861 single nuclei (Fig. 4a, Extended Data Fig. 10). We generated 10X multiome libraries from three prenatal and five postnatal brains, followed by dimensionality reduction and cell type annotation using Seurat [57] and CNV calling using epiAneufinder [58]. We used snRNAseq to determine cell types and generate gene module scores for each gene set, based on the Gene Ontology database. Unlike CNV calling from scWGS, CNV calling from snATAC lacks sufficient resolution for signature decomposition of CNVs, so we segregated nuclei broadly into “CNV neurons” vs “non-CNV neurons.” To identify CNV neurons, we inspected the distribution of the proportion of genomic regions affected by CNVs and, using Tukey’s rule, classified outlier cells as CNV neurons (Extended Data Fig. 10a-b). The correlation of CNV neuron proportion from snATAC to widespread CNV neurons determined by scWGS was high (r=0.95) between 3 donors for which both technologies were applied (Extended Data Fig. 10c).

**Fig. 4:**
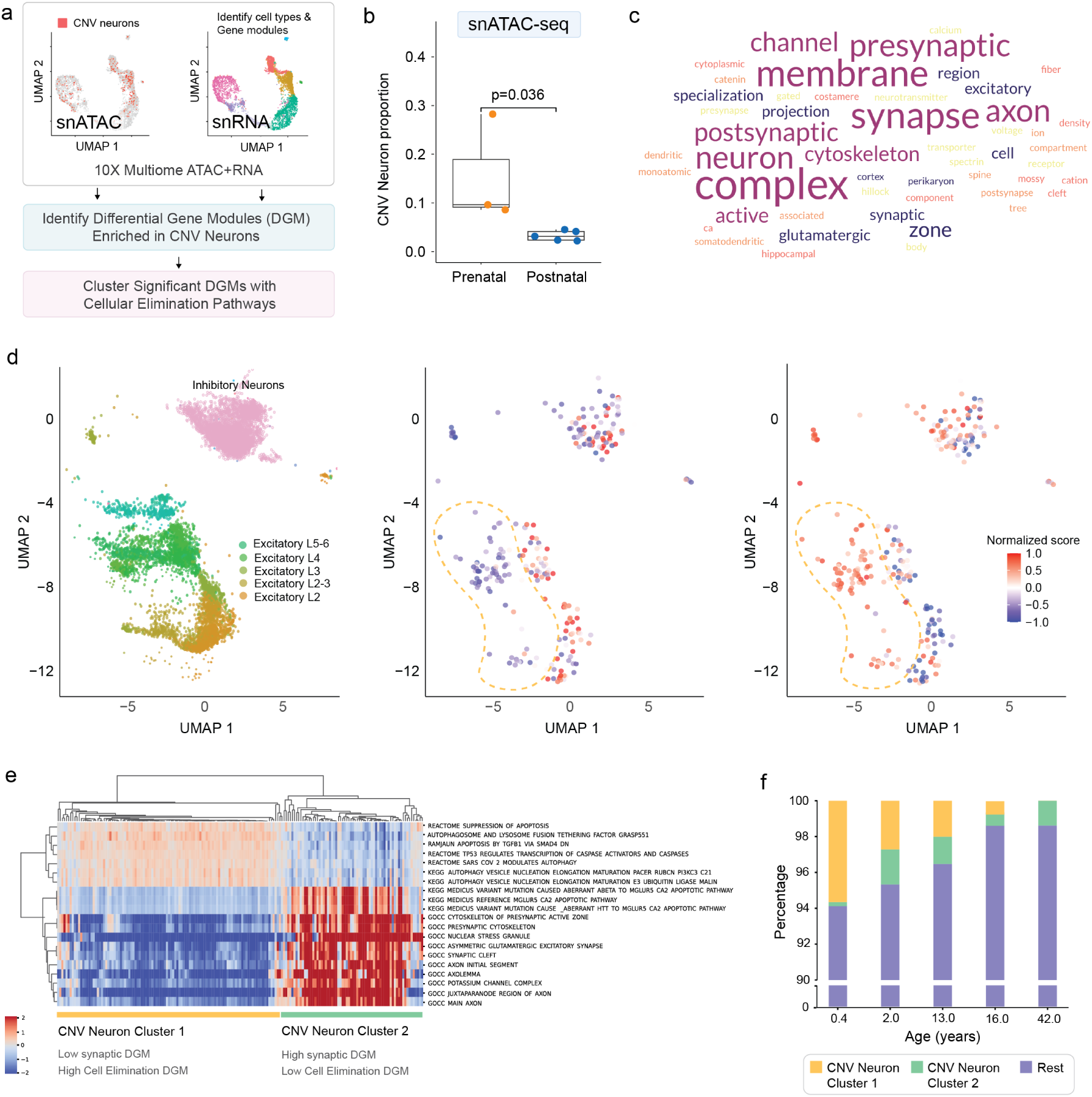
Differential gene modules (DGMs) regulating the synapse are associated with CNV neurons. **a.** Schematic of snATAC/RNA-seq workflow to identify differential gene modules (DGMs) associated with CNV states. **b.** Proportion of CNV neurons based on snATAC data in prenatal (n=3) and postnatal (n=5) brain samples show a sharp decrease after birth. Box plots indicate the median, first and third quartiles (hinges) and the most extreme data points no farther than 1.5X IQR from the hinge (whiskers). **c.** Word cloud of statistically significant gene ontology DGMs enriched in CNV neurons. **d.** UMAP visualization of snRNA-seq data from all postnatal brain cells. Left panel: excitatory and inhibitory neuron clusters. Right panels: expression of representative DGMs for synaptic function and cell death. Yellow dashed line highlights an extended cluster of excitatory neurons that show both decreased synapse gene module scores and increased cell death gene module scores (Cluster 1 neurons). **e.** Hierarchical clustering of CNV neurons based on significant DGM and cell elimination pathway expression patterns reveals two distinct clusters. Cluster 1 neurons (left) represent cells with decreased synaptic gene expression and high expression of cell death pathways, while Cluster 2 neurons (right) show the opposite pattern. **f.** Cluster 1 CNV neurons are more prevalent in younger individuals.

Our findings from snATAC confirmed a significant reduction of CNV neurons between prenatal and postnatal ages (Fig. 4b), suggesting that these CNV neurons identified by snATAC are most similar to the widespread CNV neurons observed in scWGS. We confirmed that CNV neurons have high cell quality metrics consistent with the rest of the dataset (Extended Data Fig. 10d). We found CNV neurons for excitatory, inhibitory, and maturing neuron subsets, as well as CNV progenitor cells, as expected (Extended Data Fig. 10e). CNV analysis among glial cells did not detect differences in CNV rate prenatally versus postnatally (Extended Data Fig. 10f; p=0.57, Wilcoxon rank sum), although we cannot draw firm conclusions due to limited prenatal glial sample size (*∼*1-2% of nuclei).

In order to determine pathways specifically dysregulated in CNV neurons that might drive elimination, we inferred gene modules that were differentially expressed (“Differential Gene Modules”, DGMs) between CNV neurons and non-CNV neurons. The significant DGMs associated with postnatal CNV neurons (Table S6) frequently represented components of synaptic function and connectivity (Fig. 4c). To ensure that this enrichment did not simply reflect that our analysis focused on neurons, we developed a null background by permuting CNV status within cell types. We determined that these synaptic DGMs distinguish a subset of excitatory neurons across all layers of the cortex, and a representative DGM is shown (Fig. 4d). In progenitor cells, “CNV progenitor cells” are significantly enriched for known gene modules related to cell cycle checkpoints, e.g., CDKN1A- and TP53-related pathways (Extended Data Fig. 10g). These results highlight differential regulation of dividing CNV progenitor cells versus postmitotic CNV neurons, with CNV neurons demonstrating significant differences in synapse function gene expression compared to non-CNV neurons.

Differential gene module analysis revealed that synaptic gene modules are significantly downregulated in CNV neurons compared to non-CNV neurons during the postnatal period, coinciding with the established timelines for neuronal elimination via programmed cell death. This process is characterized by extensive neuronal elimination around birth, followed by ongoing cell death postnatally through processes thought to foster appropriate synaptic connectivity during brain development [28, 32, 59–61]. Hierarchical clustering of CNV neurons based on synaptic and cell death gene modules revealed two distinct subpopulations: one subset (hereafter, “Cluster 1”) exhibited reduced synaptic module expression and upregulated cell death pathways (apoptosis and autophagy) (Fig. 4d-e), while the remaining neurons (“Cluster 2”) maintained normal to high synaptic module expression with low cell death pathway activation. The proportion of CNV neurons in Cluster 1 was highest in infant and youth samples and declined with age (Fig. 4f), consistent with the timeline of synaptic connectivity and pruning during early life [28, 33, 34]. Notably, Cluster 1 CNV neurons were absent in our midgestation samples, and limitations in third-trimester sample availability prevented assessment of late gestation. These findings suggest that Cluster 1 gene expression patterns emerge sometime during the perinatal period, and that the processes of synaptic function and cell death contribute to the decline in CNVs observed in the postnatal period.

### Genetic selection at key neurodevelopmental loci

Since the elimination of neurons across perinatal and postnatal ages favors survival of euploid neurons, we sought to identify genetic loci that are preferentially preserved or disrupted by neuronal CNVs, in order to identify signals of genetic selection. Using our scWGS data, we identified regions of the genome that were “coldspots,” regions rarely or never disrupted in surviving neurons–indicating negative selection as disruption of these regions is poorly tolerated – or “hotspots,” regions recurrently disrupted in surviving neurons that imply CNVs at these loci are tolerated and thus not negatively selected (Fig. 5a). To maximize analytical power, we included breakpoints from CNVs of all sizes. “Coldspots” frequently overlapped regions with higher gene density (Fig. 5b), a pattern not observed for “hotspots.”

**Fig. 5:**
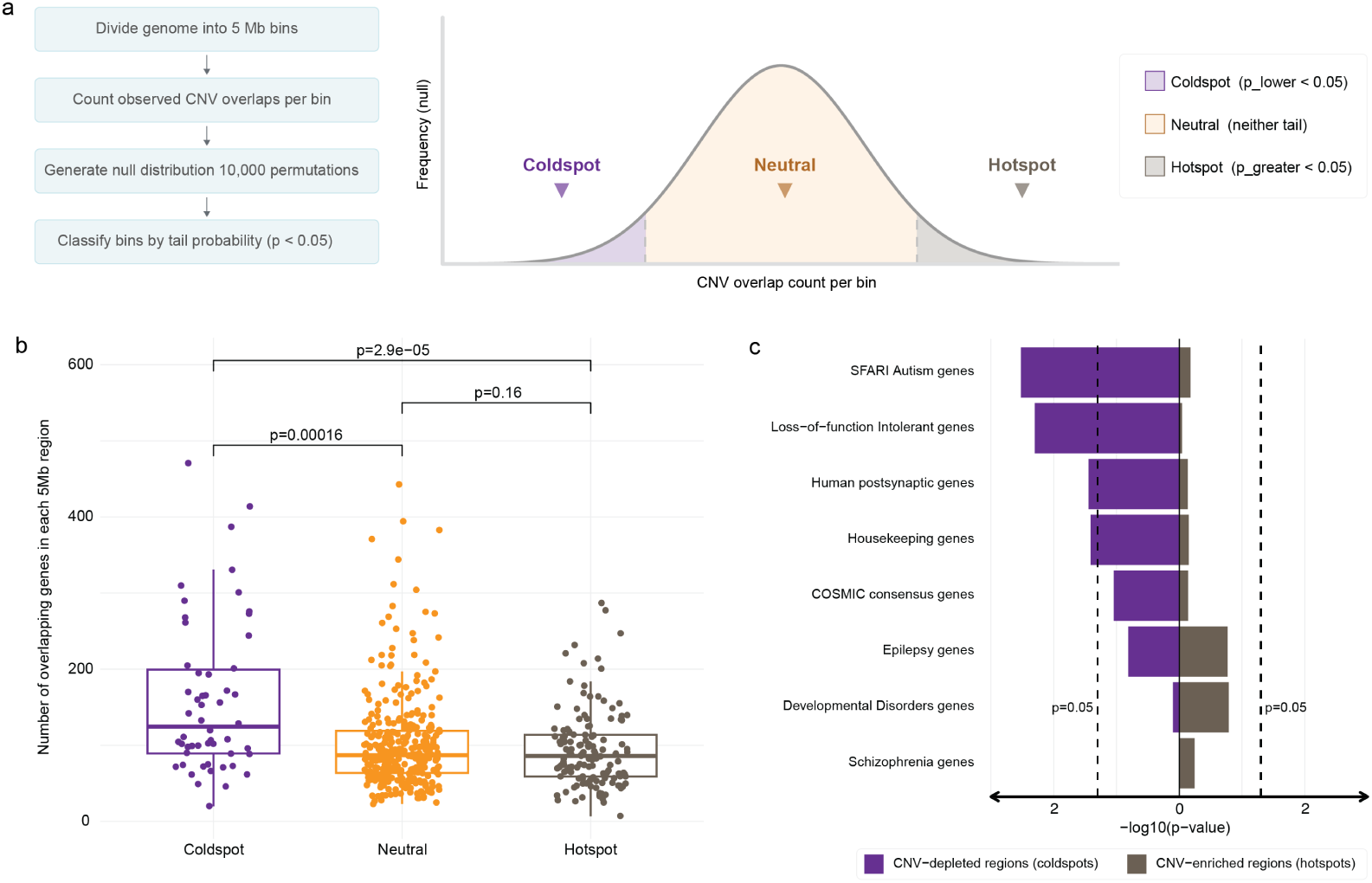
Gene-level selection of CNV. **a.** Schematic of determination of genomic hotspots and coldspots. **b.** Box plot showing the number of overlapping genes per 5Mb genomic bin classified as CNV coldspots, neutral, and hotspots (left to right). Each dot represents an individual genomic bin, with box plots showing median, interquartile range, and whiskers extending to 1.5X the interquartile range. **c.** Bar graph illustrating the enrichment p-value of specific gene categories in CNV coldspots in surviving neurons, suggesting that loss of specific gene categories may not be tolerated by the individual neurons. The dashed line indicates a significance threshold at p=0.05.

Given this observation, we hypothesized that “coldspots” may harbor key neurodevelopmental genes that neurons or progenitors may not tolerate losing. We assessed “coldspots” for enrichment of autism (SFARI [62]), epilepsy (OMIM [63]), neurodevelopmental disorders [64], and schizophrenia [64] gene sets (Fig. 5c), and found significant enrichment for Autism genes, which frequently regulate synapse formation, maintenance, and pruning [65, 66]. Further analysis using a proteomic dataset of high-confidence postsynaptic genes [67] revealed that these synaptic proteins were also significantly enriched within “coldspots” (Fig. 5c), reinforcing the critical role of synaptic genes in neuronal genome selection. “Coldspots” were also enriched for loss-of-function intolerant genes (gnomAD [68] pLI *>* 0.9) and for housekeeping genes critical to intrinsic cell function and showed no enrichment of cancer genes (COSMIC Consensus [69]). Collectively, these findings suggest that negative selection against CNV in autism-related and synapse-associated genes shapes the neuronal genetic landscape. Coupled with our DGM analyses, these data imply that regulation or disruption of synaptic gene modules—whether via largescale genomic alterations or through gene-specific loss of synaptic proteins—critically influences neuronal survival during neurodevelopment.

## Discussion

Our data suggest that widespread CNVs are a prominent feature of neurons during prenatal human neurodevelopment and are strikingly dynamic over time. These genetic abnormalities share characteristics with those seen in early post-fertilization zygotes, including signatures of genetic stress and the formation of micronuclei [6, 7, 54]. We describe two distinct phases in the elimination of neurons with CNVs: a dramatic reduction of widespread CNV neurons during the perinatal period, and a slower ongoing removal of cells that harbor CNVs over the lifespan (Fig. 6). The earlier, widespread elimination of aberrant neurons parallels the timing of neuronal cell death described during cerebral cortex development [28, 70], while our data also confirm earlier reports of slow, ongoing preferential elimination of neurons carrying subchromosomal CNVs through postnatal life and into adulthood [17, 20, 21, 70] (Fig. 6).

**Fig. 6:**
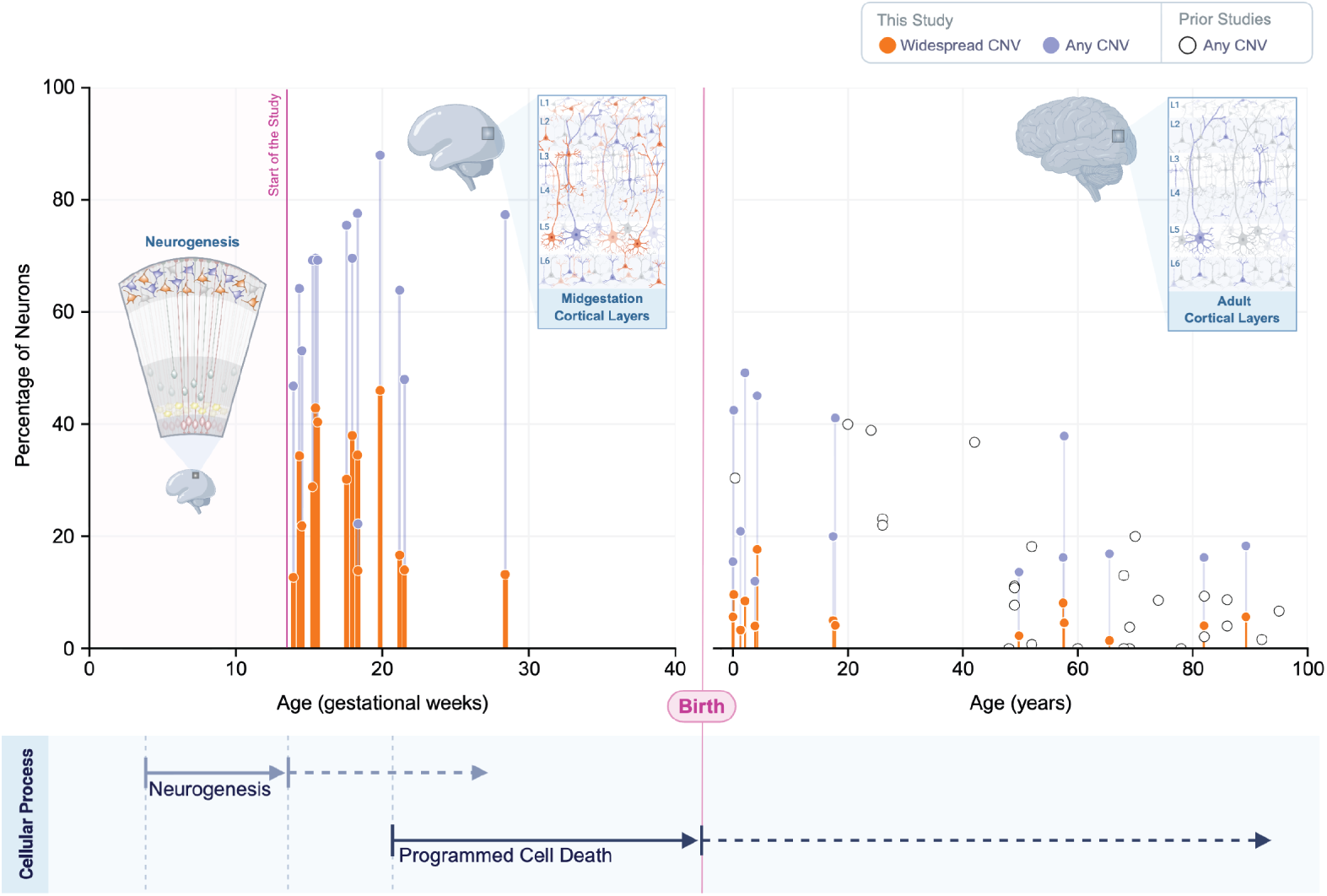
Schematic of findings from this study. Summary of CNV abnormalities across all ages for our study and prior studies of postnatal neurons (See Table S3). Neurogenesis in the cerebral cortex begins roughly at 10-12 gestational weeks, continuing to about 24 gestational weeks for excitatory neurons of the cerebral cortex and potentially until after birth for inhibitory neurons of neocortex, and produces neurons with aberrant genomes. Estimated trends programmed cell death [28, 86] establishes a large wave of cell death (∼30%) near birth, a time of extensive neuronal connectivity and maturation, followed by ongoing elimination of neurons that do not form appropriate synaptic connections. Created with BioRender.com.

Our data indicate that previously described neurobiological processes of cellular apoptosis and programmed cell death [28] may represent a core mechanism of genetic quality control for postmitotic neurons. We show that the dysregulation of synaptic gene modules in a subset of neurons with high CNV burden correlates with cellular elimination, along with selective pressure against dosage changes in critical neurodevelopmental genes related to synaptogenesis. Correct synaptic connectivity is critical in the human cerebral cortex for cell survival [71], and competition among neurons for synaptic space is known to strongly modulate neuronal cell death [72, 73]. We show that neurons with high CNV burden show dysregulated expression of synaptic pathway genes and negative selection against neurons with disruption of autism-related genes and postsynaptic genes. The force of selection is likely related to the extent of CNVs and their impact on synaptic function, with widespread CNVs predicted to impact cellular fitness more severely than smaller CNVs. The perinatal timing of the reduction of prenatal neurons with widespread CNVs proportionally mirrors the time course of a large wave of neuronal loss of *∼*30% occurring during late gestation [72, 74–76], a period of critical neuronal circuit formation, though patterns of cell death in humans have not been characterized as thoroughly as in model mammalian species due to the limited availability of appropriate tissue. Our meta-analysis of the slow postnatal decline in neurons with CNV estimates 13% neuronal loss over the course of 60 years (Extended Data Fig. 7b), on par with the estimated *∼*10% decline in aging neuron number as estimated from stereological methods [77].

Our findings of widespread CNV prenatally in noncycling neurons of the human brain reveal extensive somatic mosaicism generated in developing neural cells. At least two distinct processes appear to shape the CNV landscape. Replication stress during neuronal progenitor divisions leaves a molecular signature: CNV breakpoints cluster at early-replicating loci, and micronuclei, the cellular byproduct of replication-induced chromosomal damage, are abundant in the fetal cortex. In parallel, recurrent copy losses at BLISS-defined double-strand break hotspots implicate NHEJ-mediated repair as a second source of genomic rearrangement, one that operates preferentially at constitutively fragile loci. The age-related clearance of micronuclear content mirrors the elimination of CNV-bearing primary nuclei, consistent with a shared fate for cells carrying replication-derived genomic damage, though precise mechanisms linking micronuclei to neuronal elimination remain to be established. Together, these observations indicate that the somatic CNV landscape of mature neurons is shaped by at least two mechanistically distinct processes operating during development.

Given that persistence of aneuploid neurons would disrupt neuronal circuit function, the perinatal period emerges as a critical developmental window ensuring that neurons surviving into adulthood are predominantly euploid. The relative absence of CNVs that encompass dosage-sensitive and synapse-related genes in surviving neurons suggests intrinsic negative selective mechanisms during pre- and postnatal central nervous system development. Disruptions to the processes of perinatal neuronal genomic selection would permit genomically aberrant neurons to persist, with potentially profound implications for brain function, so it will be important to determine whether genetic factors that cause developmental disabilities show abnormalities of these selective processes. Finally, aging and neurodegenerative conditions create large burdens of other types of somatic mutations (e.g., SNVs and insertion-deletion events) with potential impacts on neuronal functions [38, 78–83], so the persistence of selective mechanisms that target neurons with damaged genomes for death may contribute to neurodegeneration later in life.

## Supporting information

Supplemental Information: Results, Methods, Tables

## Supplementary information

Supplemental materials accompanying this paper include Supplemental Information.pdf which includes Supplementary Methods and Supplementary Tables S1-S6.

## Acknowledgements

We acknowledge the sacrifices of the donors who provided tissue for this study to advance our understanding of the brain. Tissue sources were provided by the National Institutes of Health (NIH) NeuroBioBank at University of Maryland Brain and Tissue Bank, Mayo Clinic Brain Bank, the Autism BrainNet Collection, and the University of Washington Brain Development Research Laboratory (BDRL). We acknowledge the Boston Children’s Hospital Flow Cytometry Core for developing and carrying out protocols to isolate single neurons. We thank Boston Children’s Hospital Bioinformatics Core for scRNAseq analysis of cortical plate dissections. We acknowledge David S. Pellman and Greg Brunette for thoughtful discussion about identifying evidence for genetic stress. We acknowledge Peter Ly for sharing of micronuclei isolation protocols and helpful discussion. We acknowledge Takeshi Nagata for supporting micronuclear analysis.

## Declarations

### Data and materials availability

The Tn5-based single cell whole-genome sequencing data will be deposited on dbGap. BLISS BED files were downloaded from https://doi.org/10.6084/m9.figshare.18530531.v2. The ENCODE replication timing track of BG02ES was downloaded from https://genome.ucsc.edu/cgi-bin/hgTrackUi?db=hg19&g=wgEncodeUwRepliSeq. The Gene Ontology Cellular Component terms (C5) and curated gene sets (C2) were downloaded from Molecular Signatures Database [84]. The house keeping gene list is part of C2 (named “HSIAO HOUSEKEEPING GENES”). The SFARI Autism gene list was downloaded from https://gene.sfari.org/. The epilepsy gene list was downloaded from https://omim.org/. The neurodevelopmental disorder gene list and schizophrenia gene list were downloaded from http://www.brainvar.org/. The cancer driver gene list was downloaded from https://cancer.sanger.ac.uk/census. The pLI gene list was downloaded from https://genome.ucsc.edu/cgi-bin/hgTrackUi?db=hg19&g=gnomadPLI. The human postsynaptic gene list [67] was downloaded from https://static-content.springer.com/esm/art%3A10.1038%2Fnn.2719/MediaObjects/41593_2011_BFnn2719_MOESM28_ESM.xls. The postnatal reference for scRNA-seq [85] was downloaded from https://console.cloud.google.com/storage/browser/neuro-dev/Processed_data. The COSMIC pan-cancer signature panel (version 3.4) was downloaded from SigProfilerAssignment GitHub.

### Code availability

We provide the analysis code as well as detailed instructions about environment set up and software installation are available as a zip file with our submission. We intend to release the analysis code as a open-source GitHub repository in the near future.

### Author contribution

D.D.S., Y.Z., P.J.P, C.A.W. conceptualized, supervised, and designed this project. S.S., M.M., C.A.W., and P.J.P. provided resources for this project. D.D.S., J.B., V.V.V., L.J.L., T.T., F.T., and Y.Z. performed analysis. Y.A., U.G., S.Z., A.J.K., X.Q., S.C., and D.D.S. performed experiments. D.D.S. and Y.Z. wrote the manuscript. All authors provided edits to the manuscript.

## Extended Data Figures

**Extended Data Fig. 1.**
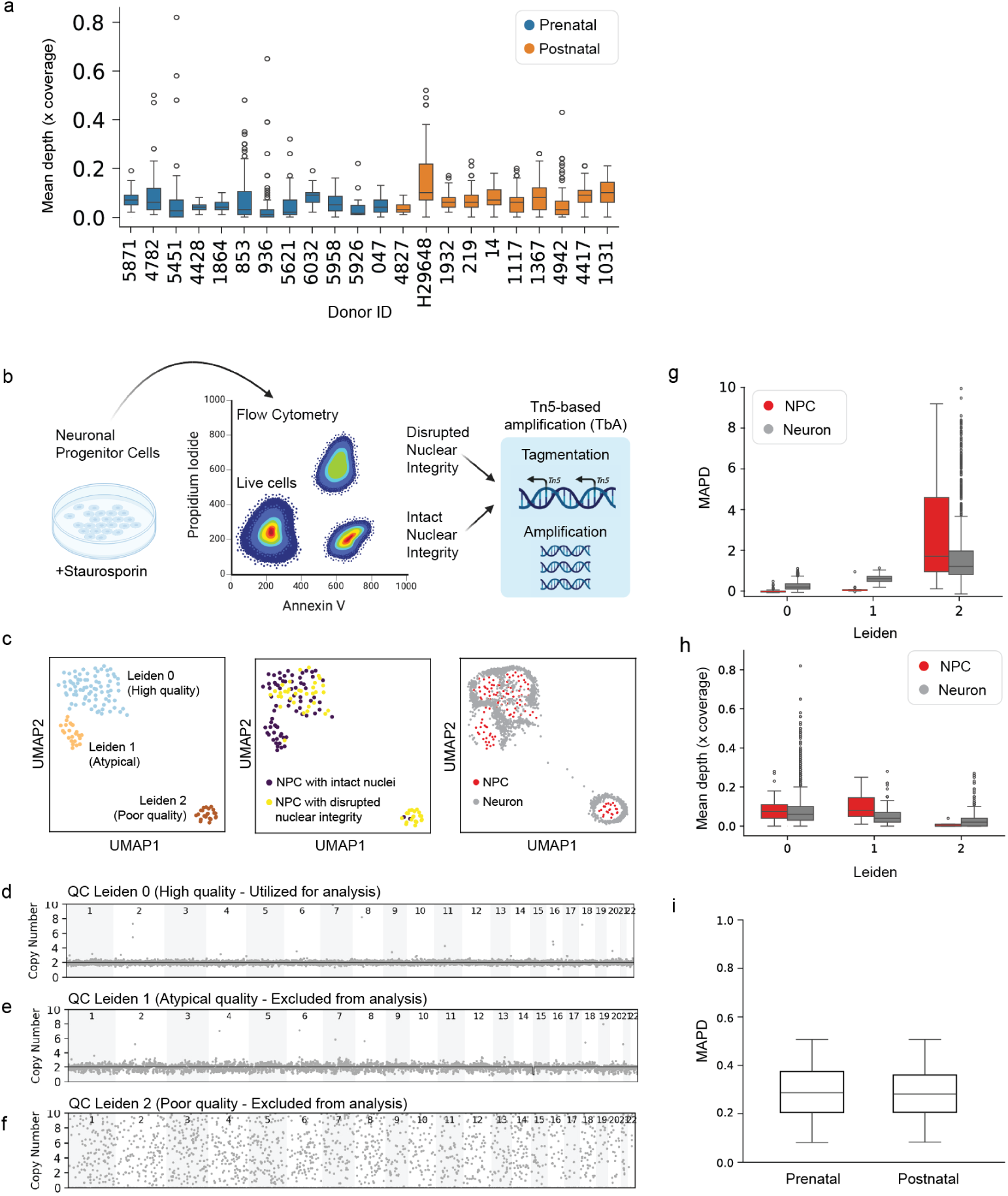
Quality control for neuronal genomes amplified by TbA. **a.** Mean coverage depth per single neuron genome amplified by TbA across human donors. **b.** Schematic of sorting neural progenitor cells for those with disrupted vs intact nuclear integrity for TbA. To form an appropriate comparison to postmortem tissue, live cells were excluded. **c.** Dimensionality reduction of binned read coverage across single-cell genomes of neural progenitor cell line (NPC) results in distinct Leiden clusters that identify high quality genomes. From left to right, UMAP colored by: (top) leiden clustering, (middle) NPC nuclei state inferred from staining, and (bottom) cell type (NPC or neuron) after label transfer. **d-f.** Representative copy number profile for QC Leiden 0-2. Only QC Leiden 0 cells were used for subsequent analyses. **g-h.** Coverage uniformity (measured by MAPD) and mean sequencing depth for NPC and single neuron from human donors from each Leiden cluster. **i.** MAPD of single neuron genomes from pre- and postnatal human donors after all QC filters have been applied.

**Extended Data Fig. 2.**
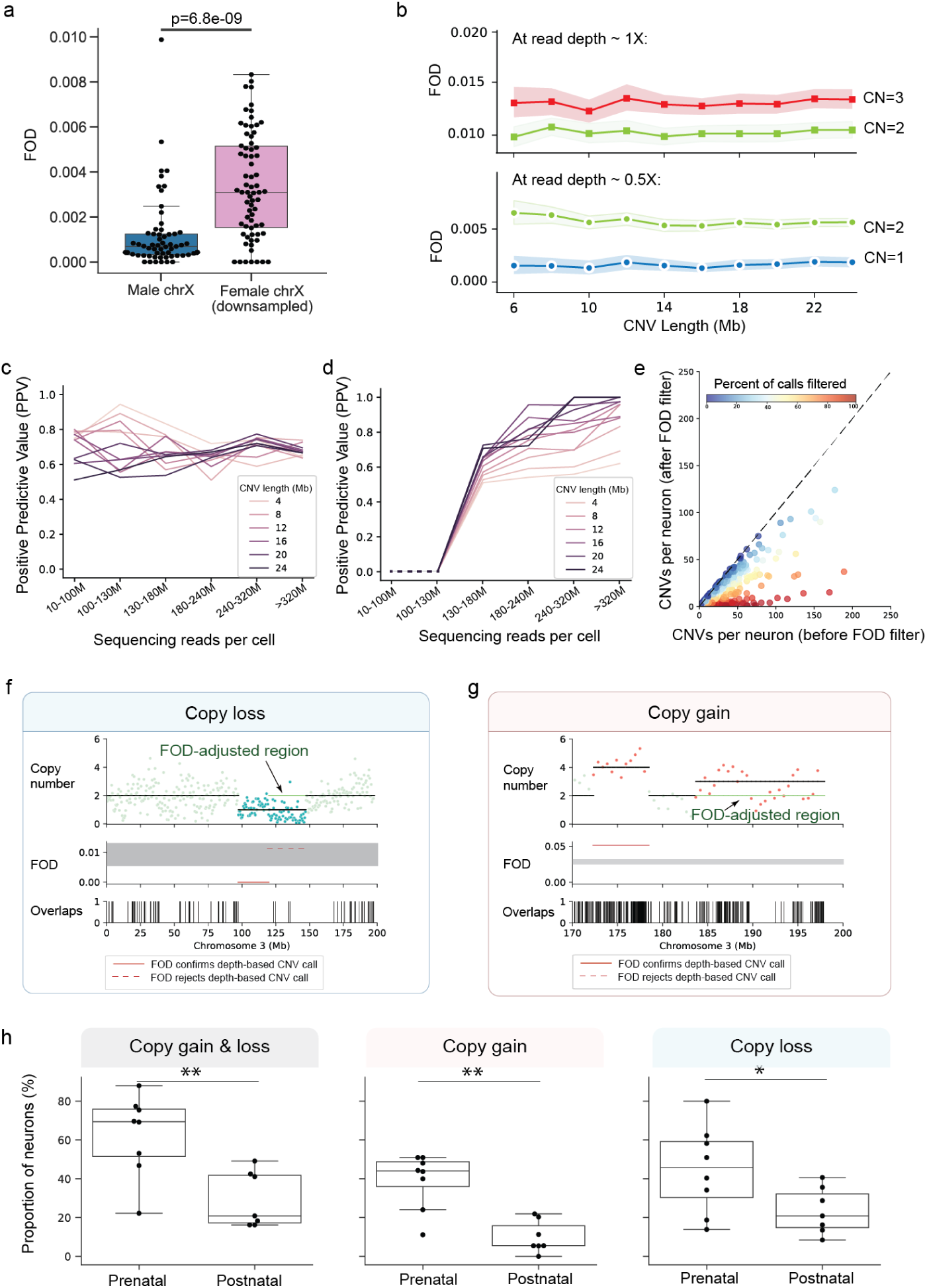
Fragment overlap density validates depth-based copy number calls. **a.** Assessment of Fragment Overlap Density (FOD) performance to distinguish copy number states at diploid (female X) and haploid (male X) chromosomes. **b.** Assessment of FOD to distinguish between single copy loss (CN=1), normal (CN=2), and single copy gain (CN=3) states using simulated CNVs by mixing defined copies of male haploid X chromosomes. Plots are separated into high and low read depth (1X and 0.5X respectively) simulations such that FOD determination of copy number cannot rely on read depth in these scenarios. **c-d.** Evaluation of FOD filter using positive predictive value in simulated single copy gains (**c**) and single copy losses (**d**) at varying sequencing depths and CNV lengths. **e.** Distribution of CNV calls per neuron before and after FOD filter, demonstrating substantial filtering of likely false positives. **f-g.** Examples of copy loss (**f**) and copy gain (**g**) from a single cell, showing copy number, FOD values, and fragment overlaps. **h.** Proportion of neurons in prenatal and postnatal samples with at least one CNV, at least one copy gain, or at least one copy loss (*p*<*0.05; **p*<*0.01.)

**Extended Data Fig. 3.**
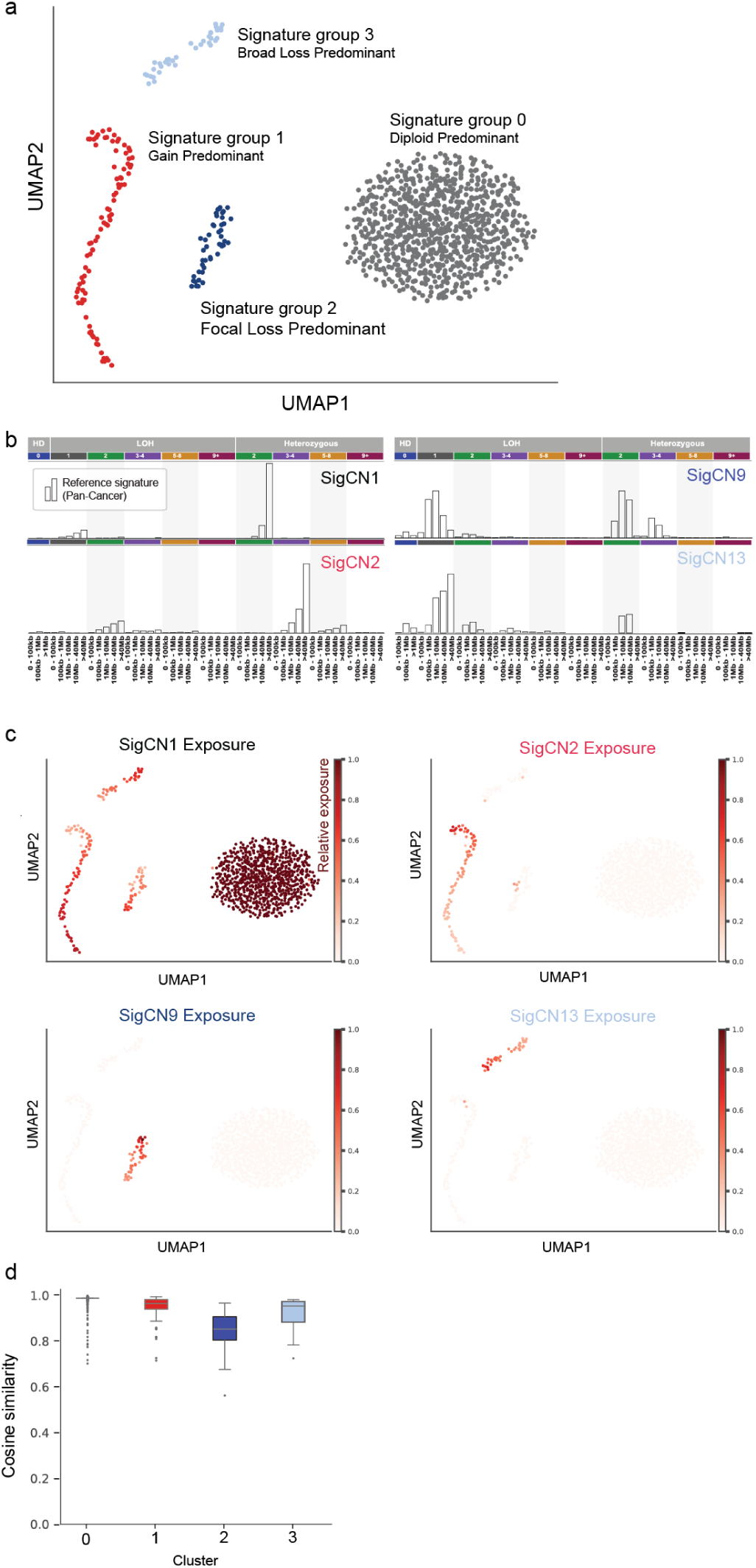
Copy number signature exposures in human neurons. **a.** UMAP of normalized exposures (i.e., relative contribution) of each COSMIC copy number signature to each neuron. We classify the neurons in Signature groups 1–3 as those with “Widespread CNV”. **b.** Reference COSMIC copy number signatures that are most prevalent in human neurons. **c.** Relative exposure of each prevalent COSMIC copy number signature in UMAP projections. **d.** Cosine similarity between composite COSMIC copy number signatures and neuronal CNV profiles.

**Extended Data Fig. 4.**
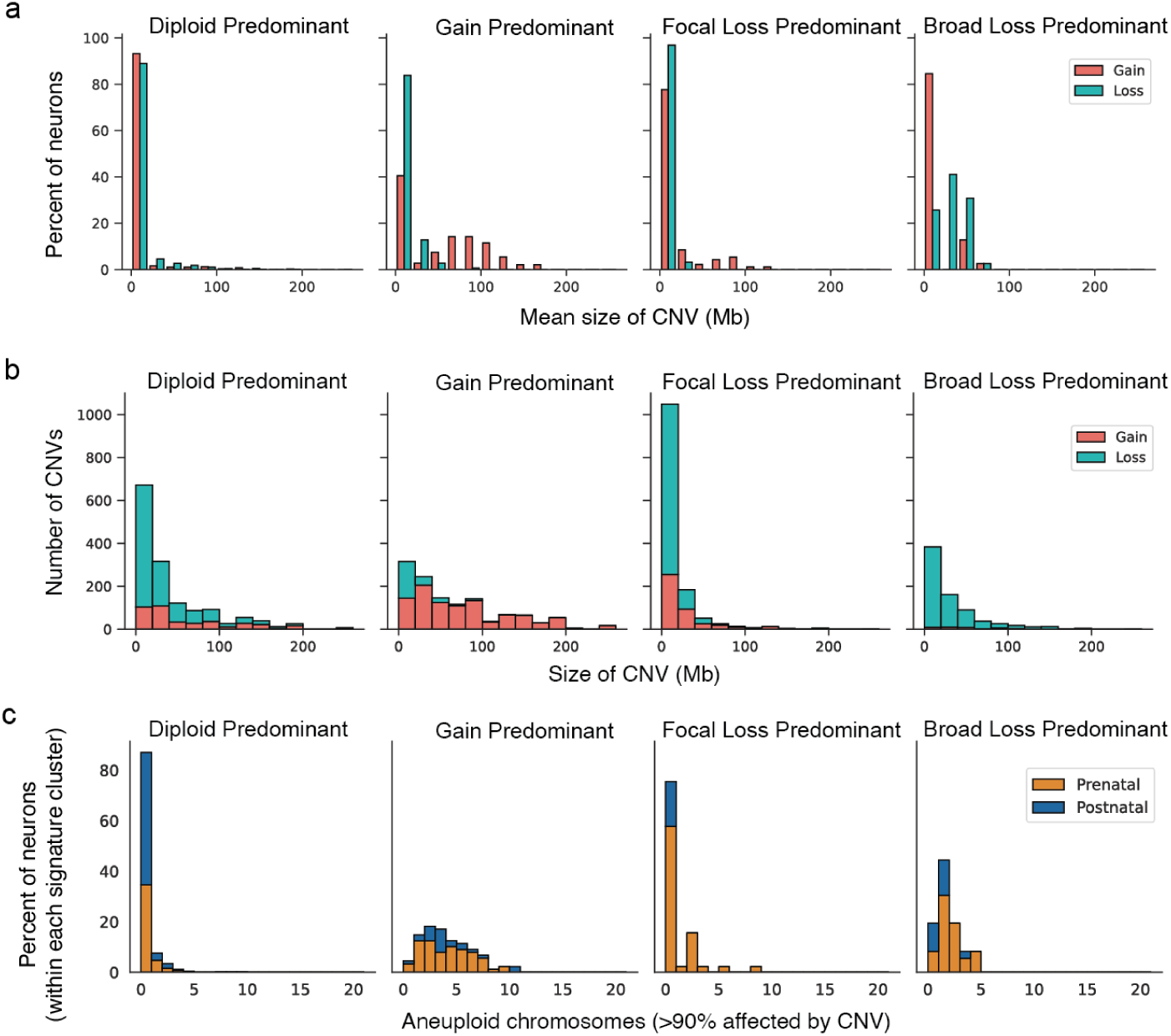
Characteristics of CNV for neurons in each COSMIC copy number signature group. **a.** Average size distribution of CNVs in each neuron signature group. **b.** Size distribution of CNVs in each signature group, accounting for only neurons that have at least 1 CNV. **c.** Distribution of aneuploid chromosomes (*>*90% affected) in each signature group.

**Extended Data Fig. 5.**
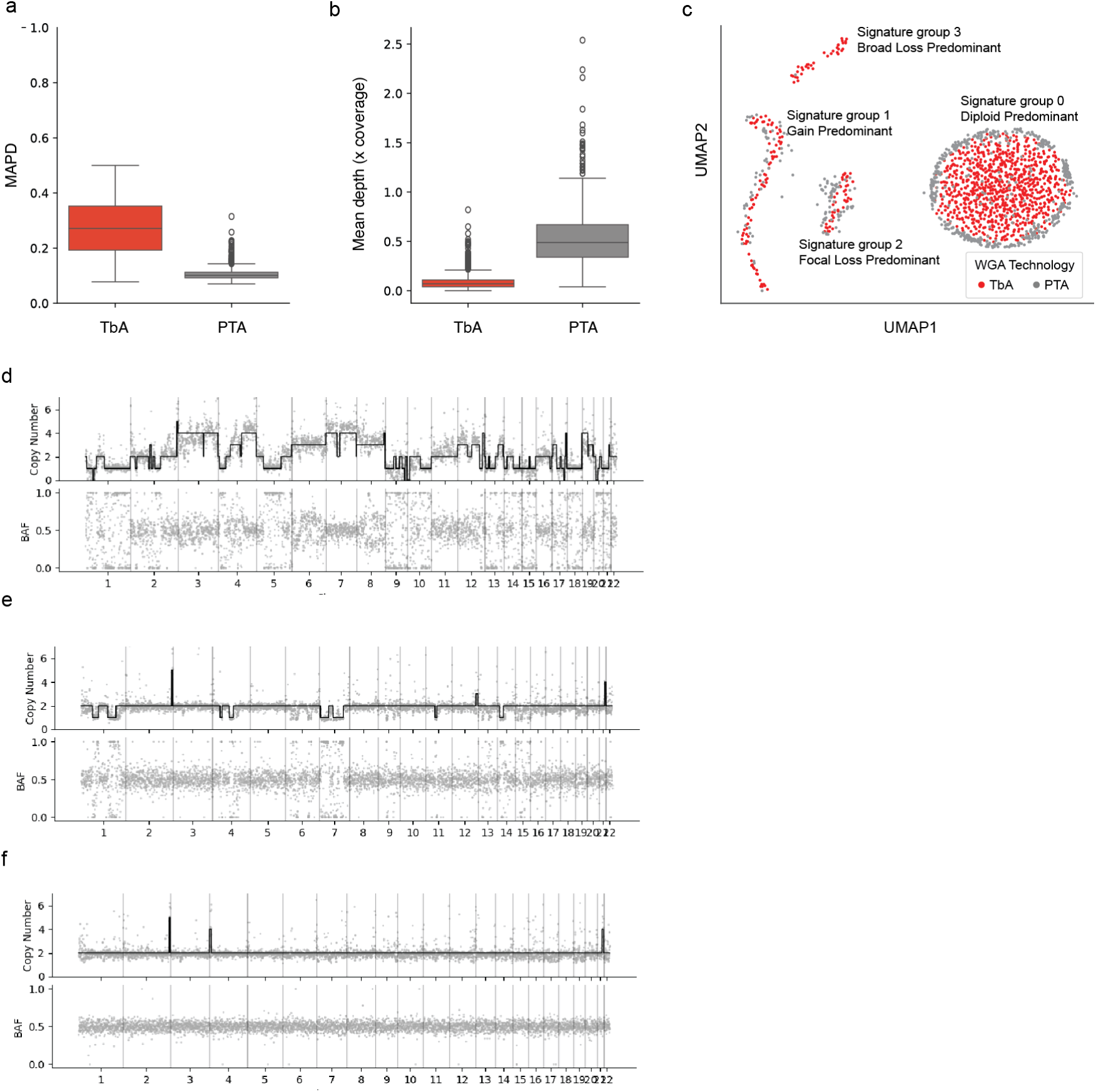
Detection of copy number signatures from primary template amplification of human single neuronal genomes. **a-b.** Coverage uniformity (measured by MAPD) and mean sequencing depth (coverage depth) for neurons amplified by PTA compared to TbA. **c.** PTA and TbA UMAP of normalized exposures for each COSMIC copy number signature shows similar signature groups. **d-f.** Representative read-depth ratio plots and associated B-allele frequency (BAF) for Gain Predominant, Focal Loss Predominant, and Diploid Predominant signature categories. CNVs at telomeres, while present in the read-depth plots, were removed from the call set due to likely telomeric artifacts for accurate copy number signature determination.

**Extended Data Fig. 6.**
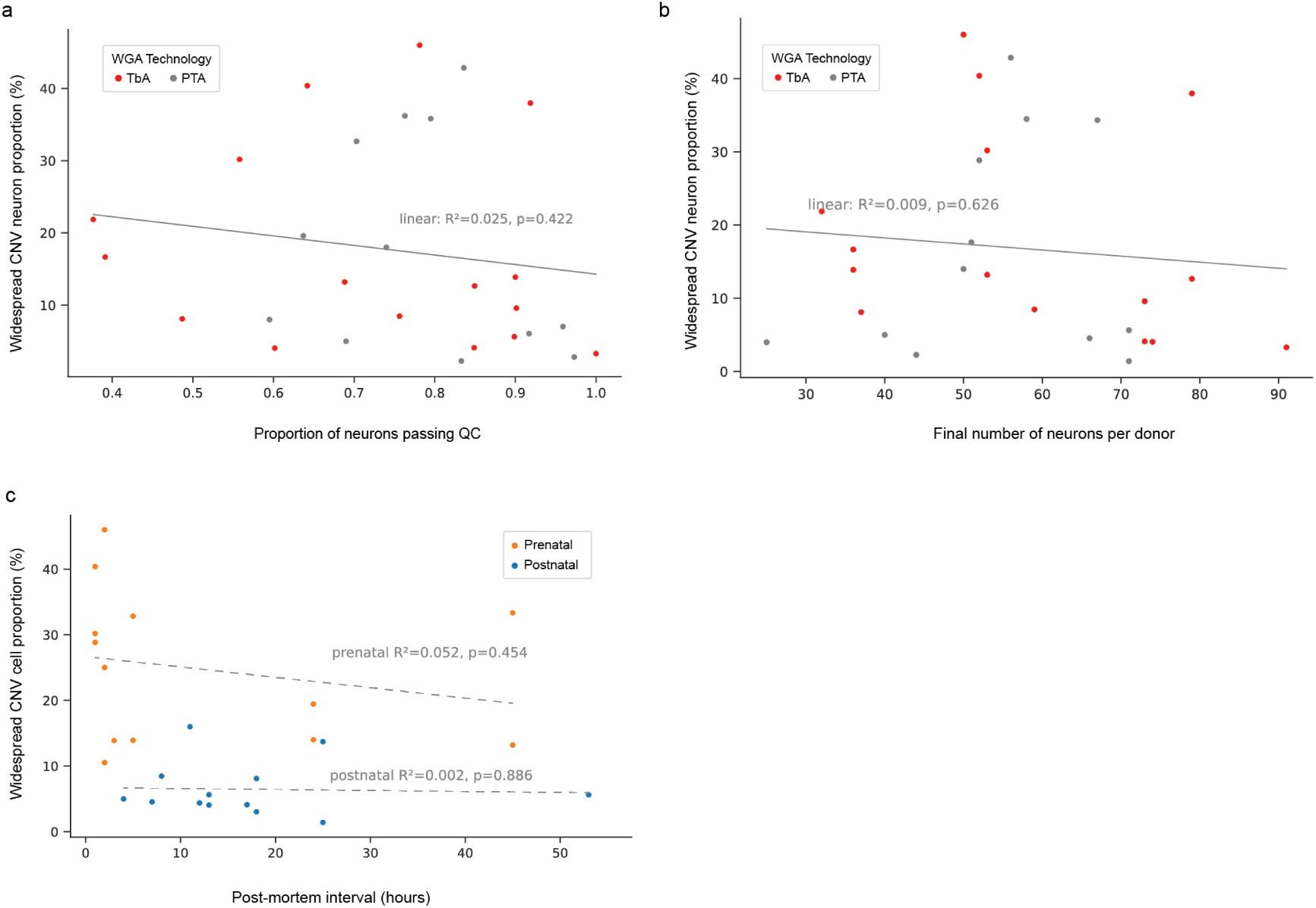
Assessment of potential confounders for the widespread CNV proportion. **a.** No significant correlation was found between sample quality (as defined by proportion of cells passing QC) and widespread CNV cell proportion. **b.** No significant correlation was found between size of final sample that passes QC and widespread CNV proportion. **c.** No significant correlation was found between postmortem interval (PMI) and widespread CNV proportion for either prenatal or postnatal groups. a-c, F-test p-value not significant.

**Extended Data Fig. 7.**
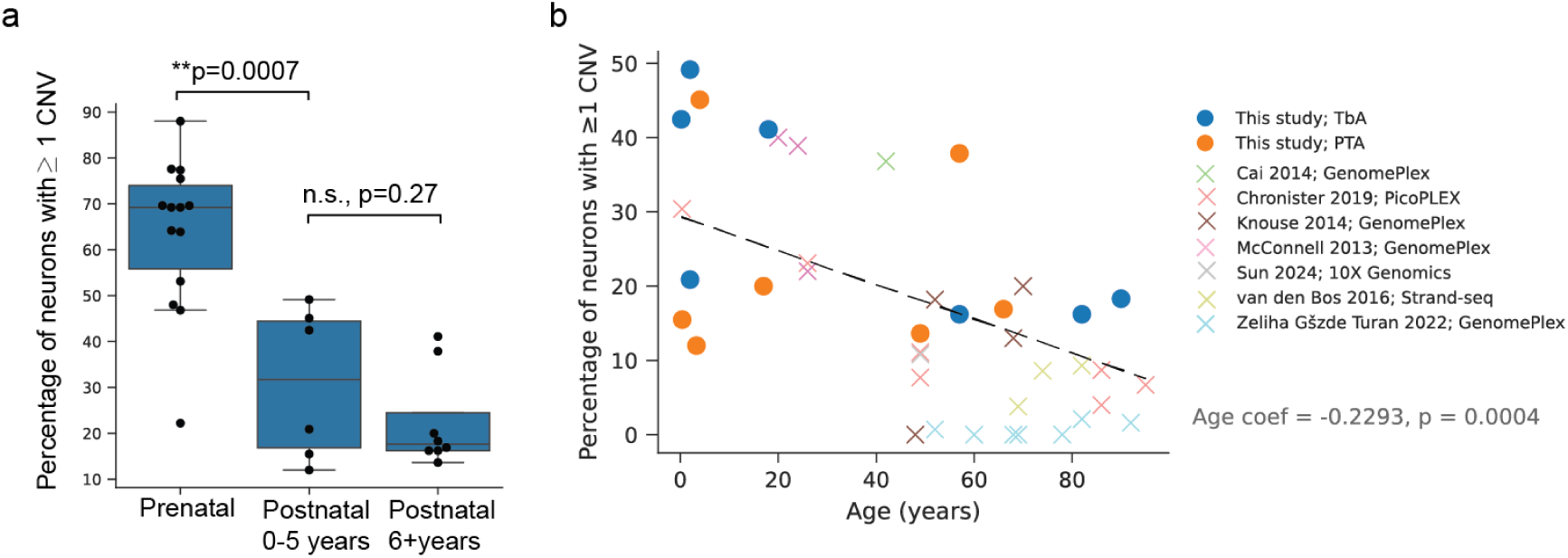
Proportion of neurons with CNV declines slowly during postnatal aging. **a.** Proportion of neurons with at least 1 CNV observed in human prenatal versus postnatal neurons. Box plots indicate the median, first and third quartiles (hinges) and the most extreme data points no farther than 1.5X IQR from the hinge (whiskers). Pvalue, Two-tailed Student t-test. **b.** Trend of postnatal neuron proportion with at least 1 CNV from our study and prior studies (Table S3). Linear mixed effects model, using each scWGA technology as covariate, was applied to obtain an estimate of annual coefficient of decrease and its p-value.

**Extended Data Fig. 8.**
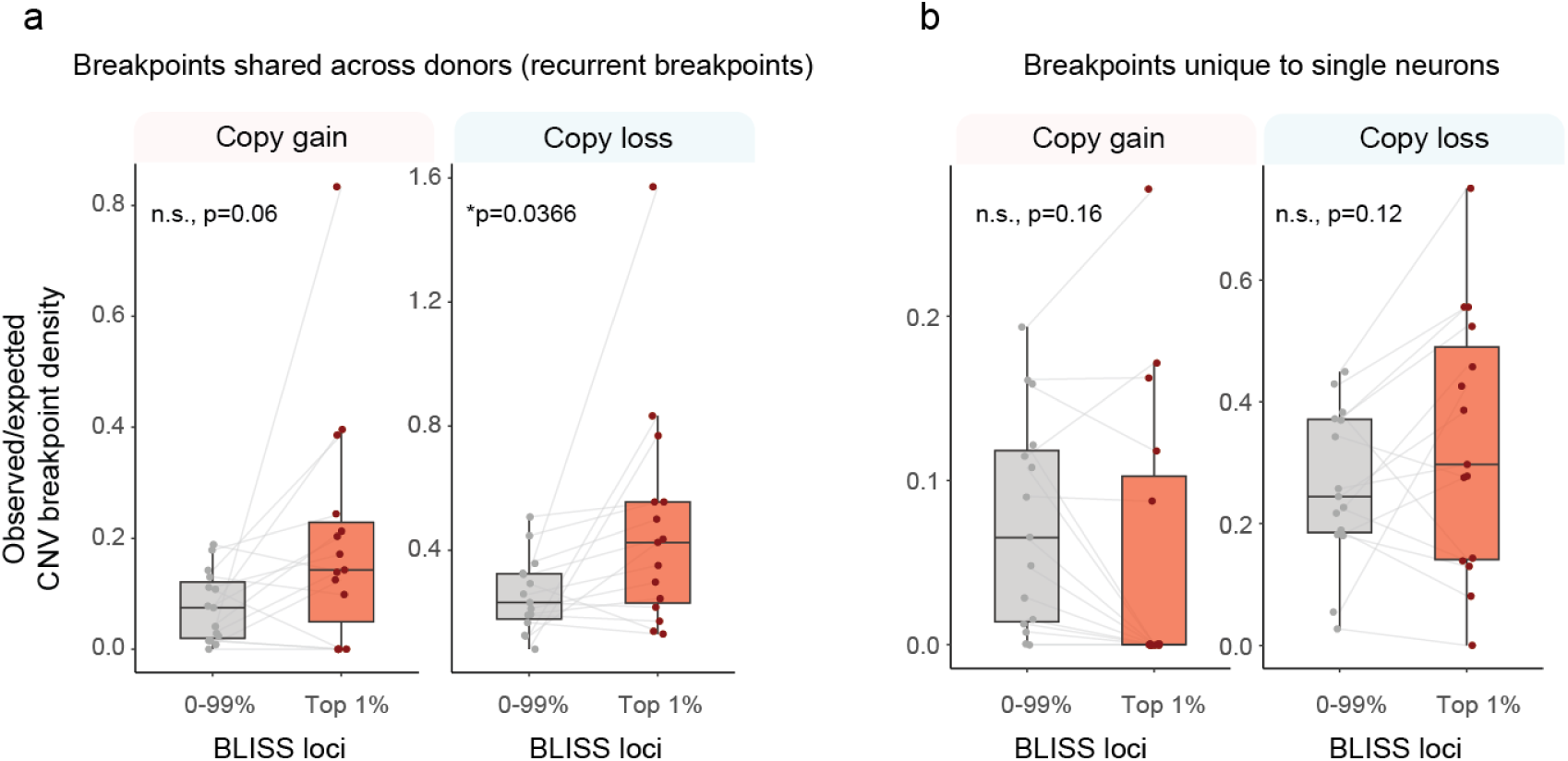
Association between CNV breakpoints and physiologic double stranded breaks detected by breaks labeling in situ and sequencing (BLISS) for CNV recurrent and shared across donors (**a**) and CNV unique to individual neurons (**b**). *p*<*0.05. n.s., not significant.

**Extended Data Fig. 9.**
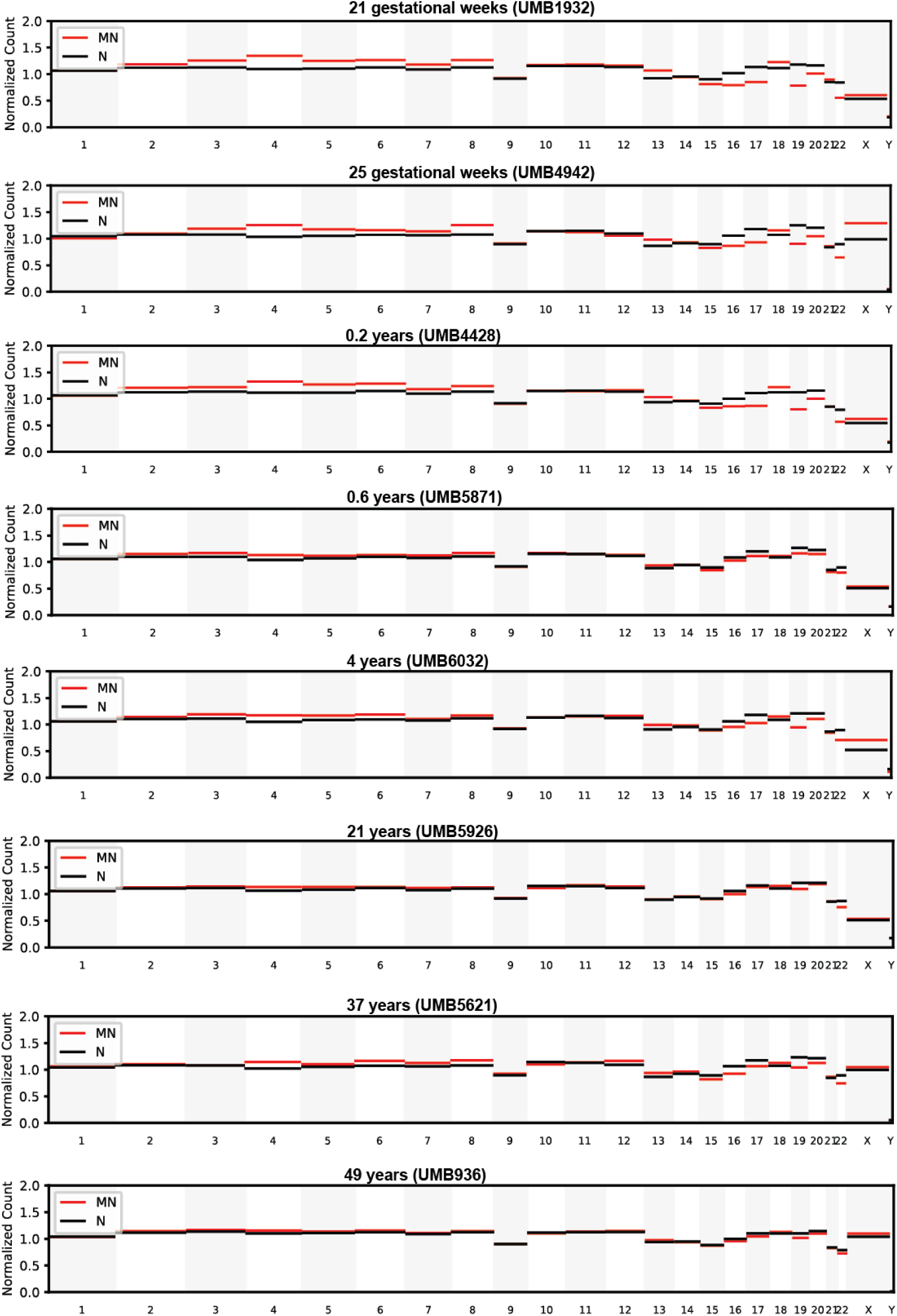
Normalized sequencing read depth across chromosomes from sucrose gradient fractions enriched for micronuclei (red) or nuclei (black), at increasing postnatal ages. With age, the depth profile of the micronuclear fraction converges toward that of the nuclear fraction, suggesting progressive clearance of micronuclei.

**Extended Data Fig. 10.**
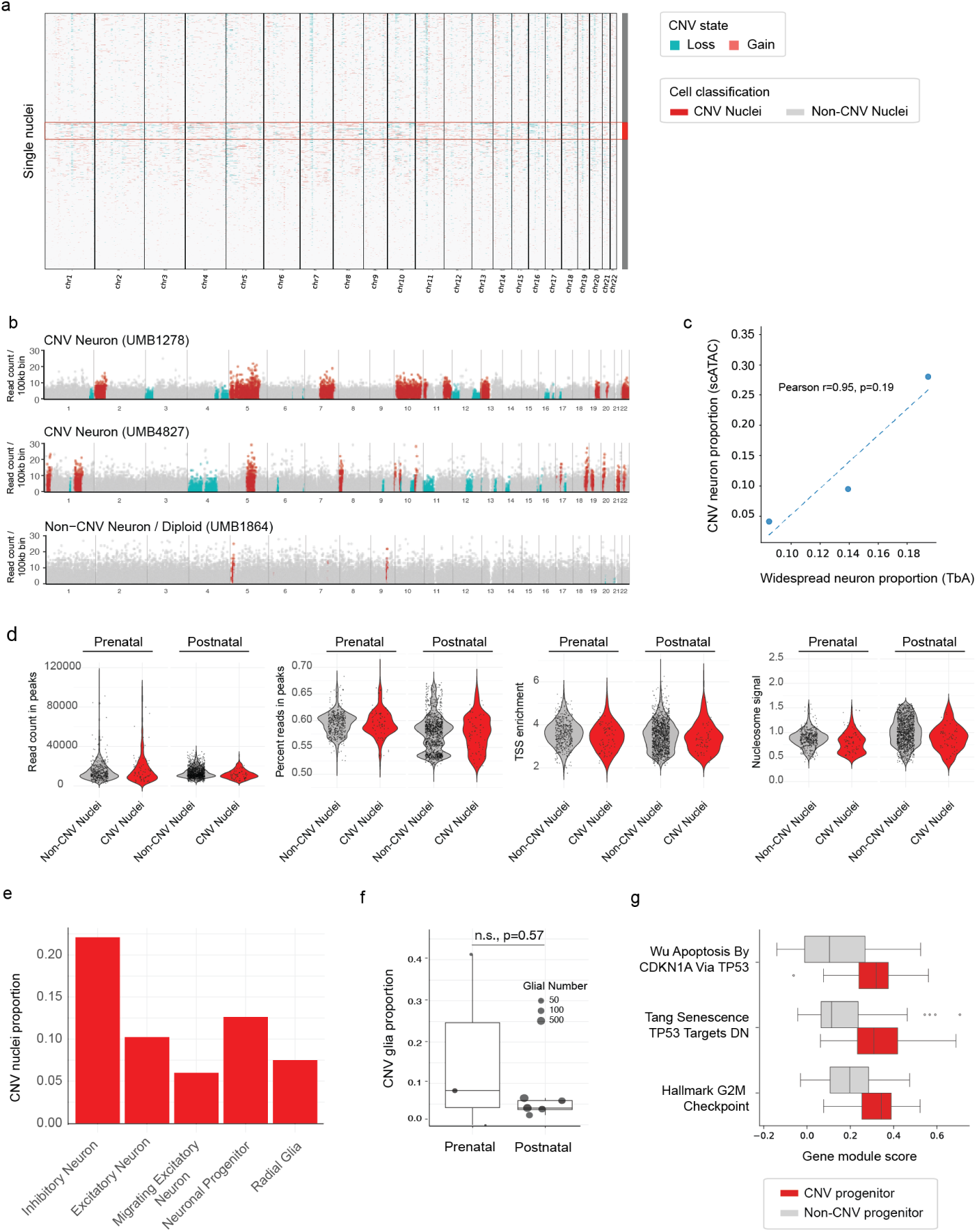
Multiomic snATAC/scRNA-sequencing identifies CNV neurons and associated gene expression patterns. **a.** EpiAneufinder CNV profiles from snATAC-seq data of a representative prenatal brain sample (H29180F4), showing chromosomal gains (red) and losses (teal). The horizontal red lines indicate a cluster of CNV neurons, cells with outlier CNV burden based on Tukey’s rule. **b.** Representative genomes of CNV neurons and non-CNV (diploid) neurons identified through snATAC-seq. Total CNV burden, rather than individual CNVs or signature, were used to identify CNV neurons due to limited specificity of individual CNV calls from snATACseq. Recurrent (likely artifactual) bins present in *>*6.17% of nuclei, based on Tukey outlier threshold, were removed. Copy gain, red. Copy loss, teal. **c.** Correlation of widespread CNV proportion (scWGS) vs CNV neuron proportion (snATAC) for three samples for which both technologies were applied. Results are correlated but not absolute due to technological limitations of CNV detection from snATAC data. **d.** Violin plots comparing quality control metrics for CNV vs non-CNV neurons at pre- and post-natal ages in snATAC-seq data. **e.** CNV cell proportions across different cell types, demonstrating CNV presence across various stages of neuronal maturity. **f.** Box plot of CNV glia proportion in pre- and postnatal brains. Note limited glial number. n.s., not significant, Wilcoxon rank sum. **g.** Box plots of DGM score distribution in CNV progenitor cells in prenatal samples at midgestation. Box plots indicate the median, first and third quartiles (hinges) and the most extreme data points no farther than 1.5X IQR from the hinge (whiskers).

## Methods

### Isolation of neuronal nuclei

Human prenatal brain tissue was obtained from the NIH Neurobiobank from the frontoparietal lobes sectioned into 40*µ*m sections with cryostat. Cortical plate was dissected using a microscalpel under a light microscope and dissociated into nuclei using gentle dounce homogenization in 1.5mL lysis buffer (10mM Tris-Hcl pH 8.0, 250mM sucrose, 25mM KCl, 5mM MgCl2, 0.1% Triton-X, 1X Compleat Mini Protease, 1mM DTT). Nuclei were pelleted by centrifugation for 10 minutes at 900G. Supernatant was removed and nuclei were resuspended in sorting buffer (1X PBS pH7.4, 0.8% BSA, 0.1*µ*g/mL DAPI). Flow cytometry was performed, gating for FSC, SSC, and DAPI content, to isolate single nuclei. For postnatal brain nuclei, staining was performed with both DAPI and Alexa Fluor 488 conjugated Anti-NeuN at 1:1000 for 1 hour at 4*^◦^*C (Sigma Aldrich MAB377X) as previously described [1].

### Single nuclei whole genome library preparation

TbA has been used in multiple contexts for single cell whole genome amplification (scWGA), under different names, with a core approach of 1) nucleosome depletion, 2) Tn5-tagmentation and direct addition of adapters, 3) library amplification. Single nuclei were lysed in microwells in 2*µ*l lysis buffer (20mM NaCl, 20mM Tris-Hcl pH 8.0, 0.15% Triton-X, 1mM EDTA, 250mM DTT, 0.05ul Thermolabile Proteinase K (NEB P8111S)) at 30*^◦^*C for 1 hour and proteinase inactivation at 55*^◦^*C for 10 minutes. Proteinase K digests histones within nucleosomes and improves accessibility of the genome by eliminating chromatin structure. Tn5 was loaded per manufacturer instructions (Diagenode C01070010-20), and 1*µ*l of Tn5 diluted 1000-fold was used per single nucleus. Tagmentation was performed using manufacturer buffers and instructions (Diagenode). Single-cell genome libraries were generated by 16 cycles of PCR using NexteraXT v2 indexes. Rows of 12 single cells were pooled for cleanup using Zymo Clean and Concentrator Kit. Each row of single cells was eluted in 42*µ*l Zymo elution buffer for subsequent Ampure XP size selection at 0.8X. Four rows (48 cells) were pooled for sequencing in equimolar concentration, and sequencing was performed on Novaseq6000 (Novogene) for a target library size of 0.5Gb per single cell library. Illumina reads were aligned to the human reference with decoy sequence GRCh37d5 (hs37d5) using bwa-mem [2], followed by duplicate marking and base quality score recalibration (BQSR) according to GATK best practices [3].

### Single-cell library quality assessment

To ensure analysis of only high quality single-cell genomes, we first removed cells with fewer than 10M total covered bases (the “total” column from GATK DepthOfCoverage, i.e. summed per-base depth across all positions), and then used neuronal progenitor cell line (NPC) data as the basis for transfer learning in order to distinguish high-quality from poor-quality sequencing profiles. The TbA libraries from single cells of a neuronal progenitor cell line (NPC) allowed us to establish a benchmark of sample quality. It is a labeled dataset of high-quality, recently viable cells as well as cells marked for apoptosis (Annexin V stain) and cells with compromised nuclear integrity through staurosporin treatment. To categorize cells computationally, we first normalized read counts for each cell in the NPC dataset using HiScanner [4]’s read-depth normalization function with bin size set to 500 kb, resulting in a cell-by-bin matrix. Next, we used Scanpy [5] to scale the read counts, standardizing variance across cells, followed by principal component analysis (PCA) for dimensionality reduction. Cell-to-cell similarity was quantified by constructing a neighborhood graph with *scanpy.pp.neighbors*. Clustering was performed using the Leiden algorithm with a resolution parameter of 0.2. After visual inspection and correlating cluster-wise read count profiles with existing replication timing tracks, library sizes, and MAPD scores, we determined that the three Leiden clusters (QC Leiden 0, 1 and 2) in NPC data are “high quality”, “atypical” and “poor quality”, respectively. We subsequently applied *scanpy.tl.ingest* to project these labels and embeddings from NPC to human brain single cell TbA data with default parameters. Only cells labeled as “high quality” (Leiden 0) and MAPD *<* 0.5 were retained. We excluded samples where the proportion of cells in Leiden 0 (per each sample batch) fell below 65%, due to concern for poor sample or batch quality. Samples in batches that yielded fewer than 25 cells after all cell-level filters were also excluded from downstream analyses.

### Single-cell CNV calling pipeline

After filtering at the cellular and sample level as above, we called CNVs on single-cell genomes. For TbA data, CNV calling was based on read depth followed by validation using fragment overlap density (FOD; Supplemental Methods). For PTA data, CNV calling jointly leveraged read depth and B-allele frequency (BAF).

#### TbA: CNV calling based on read depth

Hiscanner [4] was applied in “rdr only” mode with specific parameters optimized for our analysis. The bin size was set to 500 kb mappable positions. The hg19/GRCh37 genome was used as reference. Lambda was set to 64, and ploidy restriction was enabled (*max wgd=1* and *restrict gamma=true*). The multisample option was disabled (*multisample=false*) to process each cell individually. Hiscanner version 1.6 was installed via pip. To ensure the reliability of detected CNVs, we implemented a stringent filtering approach: CNV candidates with a p-value ≥ 0.05 or those spanning less than 5 Mb were excluded. We also removed candidates within the pseudoautosomal regions (PARs) of chromosomes X and Y, as defined in the hg19/GRCh37 genome assembly.

#### TbA: CNV validation based on FOD

To refine CNV calls, we first define a cell-specific diploid FOD distribution using the five longest segments with predicted copy number 2 (assuming no whole-genome doubling), from which the mean and standard deviation are derived. FOD Z-scores for each CNV candidate are then computed against this baseline (see Supplementary Information). Calls were retained if the FOD Z-score exceeded 1.96 for gains (CN *>* 2) or fell below −1.96 for losses (CN *<* 2); cells with a diploid mean FOD of zero were excluded. Calls failing validation retained the diploid background value of 2.

#### PTA: CNV calling based on read depth and B-allele frequency

Matched bulk whole-genome sequencing data were processed using the same alignment and pre-processing pipeline. For PTA data, Hiscanner [4] was applied in BAF+RDR mode (*rdr only=false*) with the matched bulk as reference, with a bin size of 500 kb mappable positions, lambda set to 64, *max wgd=1*, ADO threshold of 0.2, and *restrict gamma=true*. CNVs were reported as allelic copy numbers (CN_A_|CN_B_), from which total copy number was derived as the sum. Post-processing filters were applied sequentially: (1) bins called as non-diploid across cells from three or more donors were treated as recurrent artifacts and reset to CN=2; (2) CNV segments spanning less than 5 Mb were excluded; (3) putative homozygous deletion calls (allelic pattern X|0) inconsistent with the observed VAF (VAF *>* 0.1) were removed; (4) extreme copy number states (CN *>* 4) were excluded; and (5) gains (CN *>* 2) with VAF_estimate_ *<* 0.2 were removed as likely artifacts. Filter regions retained the diploid background value of 2.

### Copy number signature analysis

To identify copy number signature exposures at the single-cell level, we used SigProfilerAssignment (version 0.1.4) in conjunction with SigProfilerMatrixGenerator (version 1.2.26) [6, 7]. First, we generated a 48-channel copy number summary using the channels defined by COSMIC. Given the lack of allele-specificity in our single-cell copy number calls, we developed a conversion scheme to transform our integer copy number states into pseudo-allele-specific states. Specifically, we mapped copy numbers into allelic states as follows: 0 to 0|0, 1 to 1|0, 2 to 1|1, 3 to 2|1, 4 to 3|1, and so on up to a maximum copy number of 11 (10|1). This approach allowed us to build the input 48-channel copy number summary vectors that capture the complexity of copy number alterations while maintaining compatibility with existing signature analysis frameworks. Next, we linearly decomposed the copy number summary into copy number signature exposures. SigProfilerAssignment was executed using default parameters, utilizing the COSMIC version 3.4 pan-cancer signature panel (CN1–22, also referred to in our paper as SigCN1–22) [8] as reference. Prior to signature clustering, we excluded three cells for which CNV calling produced no callable output and cells for which SigProfilerAssignment could not produce a valid signature decomposition.

To cluster cells by signature exposure profile, we first normalized each cell’s signature exposure vector to sum to one, then added a small amount of Gaussian noise (*σ* = 10*^−^*^4^) to break degeneracies in the sparse exposure space. For TbA cells passing quality filters (MAPD *<* 0.5), we constructed a *k*-nearest-neighbor graph (*k* = 30, cosine distance) on these perturbed exposures using Scanpy [5], computed a two-dimensional UMAP embedding, and applied the Leiden community detection algorithm (resolution = 0.1) to identify four discrete groups of cells: Cluster 0 (Diploid Predominant), Cluster 1 (Gain Predominant), Cluster 2 (Focal Loss Predominant), and Cluster 3 (Broad Loss Predominant). PTA cells (after the same noise perturbation) were subsequently assigned to clusters by mapping onto the TbA reference embedding using Scanpy’s ingest function, which propagates cluster labels via *k*-nearest-neighbor label transfer in the reference UMAP space. To assess the robustness of the signature decomposition, we computed for each cell the cosine similarity between its observed copy number profile and the profile reconstructed from its assigned signature weights. A cosine similarity close to 1 indicates that the assigned signatures faithfully recapitulate the cell’s copy number landscape, whereas lower values flag cells whose profiles are poorly explained by the reference panel. Cosine similarities were aggregated and compared across clusters, serving as a per-cluster measure of decomposition confidence.

### Breakpoint distribution analysis

To investigate the relationship between CNV breakpoints and genomic features, we analyzed replication timing and DNA double-strand break (DSB) distributions. For replication timing analysis, we utilized the replication timing track of BG02ES (human embryonic stem cell) line generated by the ENCODE project [9, 10] downloaded from the UCSC Genome Browser [11]. The average replication timing signal was calculated for each 500 kb mappable bin and normalized between 0 and 1. For DSB analysis, BLISS coordinates were downloaded from FigShare [12, 13], and DSB BED files for each replicate were intersected with the 500 kb genomic bins to quantify DSB signals, which were then normalized to the per-track maximum.

To establish a background distribution for CNV enrichment analysis, 10,000 random CNVs were generated for each original CNV, matching their length distribution within autosomes (chr1-22). Peri-centromeric and telomeric regions were excluded to minimize biases, with centromeric regions defined by extending UCSC cytoband coordinates by ±5 Mb and telomeric regions defined as the first and last 5 Mb of each chromosome. Replication timing data were divided into deciles representing quantiles across the genome. For each decile, observed-to-expected (O/E) ratio was calculated by normalizing observed breakpoints (i.e., start and end sites of a CNV) to the expected number based on random simulations. Correlation between replication timing O/E ratios for CNVs from different categories (age group, signature group) was calculated, and statistical significance was evaluated using Pearson correlation tests.

To analyze the relationship between CNV breakpoints and DSBs, we compared CNV breakpoint distributions to BLISS data from six neural cell preparations: two biological replicates each from human neuroepithelial stem cells (NES), neural progenitor cells (NPC), and post-mitotic neurons (NEU). BLISS read counts were summed into 500 kb genomic bins and normalized to the per-track maximum. DSB hotspots were defined independently for each track as bins exceeding the 99th percentile of normalized signal; a bin was classified as a hotspot if it met this threshold in any of the six tracks. For each CNV, ten length-matched random CNVs were generated by placing segments of identical length at uniformly sampled positions across autosomes, and breakpoint positions (start and end of each CNV) were extracted for both observed and simulated sets. Observed-to-expected (O/E) ratios were calculated per individual as the count of breakpoints overlapping hotspot or non-hotspot bins divided by the mean count across the ten simulated replicates. NES replicates were excluded from the primary analysis due to their strong correlation with replication timing (Pearson r = 0.50–0.53, FDR-corrected p *<* 0.05), which is substantially higher than the correlations observed for NPC (r = 0.17–0.23) and NEU (r = 0.15–0.24); NPC and NEU tracks were therefore used for the aggregated hotspot analysis. O/E ratios were capped at 4 for visualization. Statistical significance was assessed using paired t-tests comparing each individual’s hotspot O/E ratio to their non-hotspot O/E ratio (n = 15 pairs each for gains and losses), with p-values adjusted for multiple testing using the Benjamini–Hochberg FDR procedure.

CNV breakpoints were further stratified by recurrence. A breakpoint was classified as recurrent if the same genomic position was observed in more than one donor; breakpoints detected in more than two donors were used for full-CNV recurrence analysis. A breakpoint was classified as private if it was detected in exactly one cell from a single donor. Within-donor clonal breakpoints—shared by two or more cells from the same individual—were analyzed separately. BLISS hotspot enrichment analysis was performed independently for each recurrence class.

### CNV gene set enrichment analysis

To identify hot and cold spots in Diploid Predominant neurons, we adapted an approach similar to Sun *et al* [14]. We defined genomic bins of 5 Mb by aggregating the existing 500 kb bin coordinate set that was used in CNV calling, with any remaining small segments at the end of the chromosome discarded. Each 5 Mb bin was assigned an empirical overlap count, representing the number of overlaps with observed CNVs. We then performed 10,000 permutations, as outlined in the “Breakpoint distribution analysis,” preserving the CNV sizes. For each permutation, the number of overlaps with random CNVs was also calculated, allowing us to generate a null distribution for each bin. P-values were calculated to assess whether the observed CNV overlaps deviated significantly from the random expectation, with separate p-values for enrichment (“hotspots”) and depletion (“coldspots”). Adjusted p-values were computed using Bonferroni correction, and significant hotspots and coldspots were identified accordingly. Each 5 Mb bin was further evaluated for overlaps with gene sets, including housekeeping genes (Gene Ontology), loss of function intolerant genes (pLI [15]*>*0.9), SFARI [16] Autism genes, neurodevelopmental disorder [17], schizophrenia [17] gene sets, post-synaptic gene sets [18], and cancer driver genes from COSMIC [19], and epilepsy (OMIM [20] symptom search “epilepsy”). Based on their adjusted p-values, bins were classified as hotspots or coldspots, using a significance cutoff of 0.05. Bins not reaching the significance threshold were classified as neutral. To assess gene set enrichment within hotspots and coldspots, hypergeometric tests were performed to evaluate over-enrichment of each gene set within these regions compared to all genomic bins.

### Single-nuclei multiomic ATAC/RNA library preparation

Nuclei were isolated without dissection as described in “Isolation of neuronal nuclei” and permeabilized after flow cytometry according to the Nuclei Isolation from Complex Tissues for Single Cell Multiome ATAC + Gene Expression Sequencing protocol from 10X Genomics (Demonstrated Protocol, CG000375). Chromium Next GEM Single Cell Multiome ATAC + Gene Expression libraries were prepared according to the manual with adaptations to also enable detection of complementary DNA strands (duplexes) [21]. Duplex information was not utilized for this study.

### Cell type and CNV annotation of multiomic snATAC/snRNA data

The multiomic dataset consists of snRNA-seq and snATAC-seq data from prenatal and postnatal brain samples, produced following the 10X Genomics protocol. Data were processed using the 10x Genomics Cell Ranger ARC pipeline (version 2.0.1). Subsequently, the dataset was analyzed with Seurat (version 5.0.1) [22] and Signac (version 1.12.0) [23] R packages.

#### QC filter of nuclei from snATAC-seq

For both prenatal and postnatal brain samples, nuclei were filtered on several metrics. First, multiplet detection was performed on snATAC-seq data using AMULET, after which cells classified as multiplets were filtered out. Next, filtering thresholds for several metrics were then determined by visual inspection of rank plots of each metric for all samples. For total RNA molecule count, RNA feature count (number of genes), and total ATAC fragment count, a lower threshold of 1000 was used to remove empty background nuclei and no upper threshold was used given multiplet removal had already been performed. Finally, for both nucleosome signal and transcription start site (TSS) enrichment values from snATAC-seq, nuclei were removed with lower (0.32 for nucleosome signal and 1.75 for TSS enrichment) and upper (1.59 for nucleosome signal and 6.80 for TSS enrichment) thresholds determined by the rank plots. Additionally, a lower threshold of 0.5 was determined from the rank plots for the fraction of reads in peaks.

#### Cell type annotation based on snRNA-seq

Cell type annotation was performed by mapping each sample to a reference sample. We chose separate references for prenatal and postnatal due to the differences in cell type composition between prenatal and postnatal cells. For the prenatal samples, 29180 F4 was chosen as reference, and for the postnatal samples, the downsampled reference was used from a prior published dataset [24]. The prenatal reference was manually annotated by first performing unsupervised clustering on normalized, PCA-transformed RNA count data. The resulting clusters were then manually annotated using known marker genes for different cell types. A UMAP (Uniform Manifold Approximation and Projection) representation was generated for the prenatal reference. For the postnatal reference, the pre-existing annotations from Herring *et al* [24], based on clustering performed in the paper, were used. The remaining non-reference “query” samples were then projected onto the reference UMAPs using Seurat’s MapQuery function. The MapQuery function transferred the manually annotated cell type labels from reference to query samples, resulting in complete cell type annotation for all samples in the multiome dataset.

#### CNV inference from snATAC-seq

CNV calling was performed using the R package epiAneufinder (version 1.0.3). [25], with a window size of 100 kb, and a minimum fragment number of 20,000 per sample. The parameter *minCNVsize* was set to 0, meaning that the minimum callable CNV size was 1 bin, i.e. 100 kb. After CNV calling was complete, the CNV calls were combined across all samples by taking the intersection of bins in all samples. For the resulting bins, for each cell across all samples, the percentage of bins with a CNV (either gain or loss) was calculated. The resulting distribution of percentage of bins affected was used to determine a cutoff for our definition of CNV Neurons or CNV Cells. Tukey’s rule was applied for identifying outliers, such that any cell with a percentage of bins greater than the third quartile plus 1.5 times the interquartile range (Q3 + 1.5IQR) was classified as an outlier. These outliers were then classified as CNV neurons, and non-outliers as non-CNV neurons.

### Differential gene module (DGM) analysis

We analyzed the snRNAseq part of single-cell multiome data separately for prenatal samples (n=3; UMB4827, UMB1932, 29180 F4) and postnatal samples (n=5; APGEQ, U1790, U4643, UMB1278, UMB1864) due to differences in cell type composition. For each age group, sample-specific Seurat objects were merged into a combined dataset while preserving sample identifiers. Gene expression values were log-normalized with a scale factor of 10,000. To assess pathway activation, we calculated module scores using gene sets from the Molecular Signatures Database [26] (MSigDB), including Gene Ontology Cellular Component terms and curated gene sets (C2). For C2, we only included gene sets that matched the keywords “senescence,” “apoptosis,” “autophagy,” and “caspase” – hereafter referred to as “cell elimination C2 pathways.” Module scores were computed using Seurat’s *AddModuleScore* function using default parameters, which calculates the average expression of each gene set in a given cell after controlling for gene expression program complexity by subtracting the aggregated expression of randomly selected control gene sets with similar average expression levels.

To test for associations between gene module scores and CNV neuron status, we implemented a stratified permutation testing approach. For each gene module, we calculated the observed difference in mean module scores between CNV and non-CNV cells (hereafter referred to as “absolute observed difference”). Statistical significance was assessed through 10,000 permutations, where CNV neuron labels were randomly shuffled within each stratum defined by sample and cell type, preserving the underlying data structure. P-values were calculated as the proportion of permuted differences that exceeded the observed difference in absolute value and were adjusted for multiple testing using Bonferroni correction. In order to avoid confounding by cell-type specific gene expression differences, this analysis was performed separately for each major cell type (excitatory neurons, inhibitory neurons, progenitors, and rest) and developmental stage (prenatal and postnatal).

### Clustering of CNV neurons based on DGM and cell death pathways

To identify distinct subpopulations of CNV neurons based on their pathway activation patterns, we performed hierarchical clustering on the normalized gene module scores from a selected set of pathways. Since clustering on all available pathways could introduce noise from non-informative or weakly differential pathways, we focused on the top 10 most differentially activated gene modules from GOCC and cell elimination C2 pathways, ranked by the absolute observed difference (see DGM analysis section above for definition of “absolute observed difference” and the selection of cell elimination C2 pathways). The module score matrix was normalized by subtracting the mean and dividing by the standard deviation across cells. Hierarchical clustering was performed using average linkage with the cosine distance metric. We determined an optimal cluster number (n=2) through visual inspection of the dendrogram and cluster stability.

To visualize pathway activation patterns, we generated heatmaps using the seaborn *clustermap* function with row clustering enabled for gene sets and column clustering for cells. The resulting visualization revealed two main clusters of cells with distinct pathway activation patterns. Cell cluster assignments were validated through multiple approaches, including assessment of cluster sizes (Cluster 1: 1,667 cells; Cluster 2: 2,487 cells) and evaluation of enrichment of CNV neurons using Fisher’s exact test. The association between cluster membership and CNV neuron status was quantified using odds ratios with 95% confidence intervals. Multiple testing correction was performed using the Bonferroni procedure.

