## Supplemental Information: Results, Methods, Tables for "Perinatal Elimination of Genetically Aberrant Neurons from Human Cerebral Cortex"

### Supplemental Results

#### Fragment Overlap Density (FOD) as a novel validation method for single-cell copy number analysis

##### Experimental design for FOD validation

Conventionally, CNV detection relies on depth-based calls validated by B-allele frequency (BAF) measurements. However, this approach becomes challenging in single cells due to amplification noise and sparse coverage. To validate HiScanner[1]-derived copy number variant (CNV) calls in single-cell whole genome sequencing (scWGS) data, we developed a novel metric called Fragment Overlap Density (FOD) that leverages the unique properties of Tn5-transposase-based amplification (TbA) library construction. FOD is defined as a ratio of overlapping fragments to total number of fragments in a region of interest, resulting from the unique properties of Tn5-transposase-based library construction in which all subsequent amplified products retain the start and end positions of the original tagmented DNA molecule. FOD varies based on the total number of alleles present in a locus in a single cell, effectively distinguishing between copy loss region where only one DNA allele is present–no fragment overlaps should be observed–versus copy neutral or copy gain region in which respectively more overlapping fragments would be observed. FOD provides additional information for CNV inference where the read-depth signal is ambiguous, and is more flexible than BAF: FOD can be utilized for datasets with as few as one cell or datasets without phased or matched bulk, as long as Tn5-transposase is used in the first step of single-cell library construction. Our FOD metric provides an orthogonal validation metric for single-cell CNV calls by quantifying the proportion of overlapping DNA fragments within a genomic region.

##### Theoretical and simulated evidence for FOD performance

We first validated FOD using simulated data constructed from male samples, where X chromosomes provide a natural control for copy number state detection (Extended Data Fig. 2a). As expected, female X chromosome (diploid) samples demonstrated significantly higher FOD values compared to male X chromosome (haploid) samples (two-sided Student's t-test p = 6.80e-09), confirming that FOD can distinguish between diploid and haploid copy number states.

To systematically evaluate FOD's performance in distinguishing copy number states across various sequencing depths and CNV sizes, we conducted simulation experiments with controlled copy number states (Extended Data Fig. 2b). Our simulations demonstrated that FOD values consistently separated different copy number states (CN=1, CN=2, CN=3) at both high (1X) and low (0.5X) sequencing depths. Importantly, this separation remained consistent across CNV lengths ranging from 6Mb to 22Mb, indicating that FOD is robust to variations in CNV size.

We further validated the FOD metric through systematic simulations of single-copy gains (Extended Data Fig. 2c) and losses (Extended Data Fig. 2d) across various sequencing depths. For single-copy gains (Extended Data Fig. 2c), our analysis revealed positive predictive values ranging from approximately 0.6-0.8 across all sequencing depth bins from 10-100M to >320M reads per cell. Notably, performance was consistently robust across all tested CNV lengths (4-24Mb), with some variability observed in the lower sequencing depth ranges (10-100M reads).

For single-copy losses (Extended Data Fig. 2d), we observed a distinct pattern where positive predictive values were near zero at low sequencing depths (10-100M reads) but sharply increased between 100-130M reads per cell. Once sequencing depth exceeded 130M reads, positive predictive values for loss detection rose dramatically to approximately 0.6-0.8 and continued to improve with increasing sequencing depth, reaching >0.9 at the highest depth bin (>320M reads). This performance pattern was consistent across all CNV length categories, indicating that while FOD requires sufficient sequencing depth to reliably detect copy losses, it performs robustly once this threshold is reached.

#### Empirical application and impact on CNV calling

When applied to our scWGS dataset, FOD validation significantly refined our initial depth-based CNV calls (Extended Data Fig. 2e). The comparison between CNV counts per neuron before and after FOD filtering revealed that FOD rejected many false-positive calls, particularly in cells with high initial CNV counts. This effect is illustrated by the deviation from the diagonal line (y=x), indicating that FOD provided substantial correction to the raw depth-based calls. We provide representative examples that demonstrate FOD's utility in both validating true CNVs and rejecting false calls: Extended Data Fig. 2f shows a copy loss event where FOD values were significantly reduced compared to the diploid background, confirming the depth-based call. Conversely, Extended Data Fig. 2g illustrates a case where FOD rejected a depth-based copy gain call due to inconsistent fragment overlap patterns.

In conclusion, our FOD metric provides a robust, independent validation method for single-cell CNV detection that complements traditional depth-based approaches. By leveraging the molecular properties of Tn5-transposase library preparation, FOD enables more accurate CNV calling in single cells without requiring matched bulk samples or phased genotypes, representing a significant methodological advancement for CNV detection in single-cell genomics.

### Supplemental Methods

#### Using fragment overlap density in TbA scWGS to improve CNV detection

In TbA scWGS, the fragment overlap characteristics can be used to improve CNV detection in two ways: amplification bias reduction and CNV candidate validation.

#### Amplification bias reduction

Tagmentation by Tn5-transposase on each DNA molecule is random and independent, creating unique sequences distinguished by read start and stop sites. Therefore, removing reads mapped to identical start and end coordinates in the genome can effectively reduce amplification bias prior to depth-based CNV calling [**?** ].

#### Informatic validation of CNV calls

FOD distinguishes between single alleles (copy number 1) which mechanistically can only have the 9-bp strict overlap between adjacent Tn5 tagmentation sites [2], and higher allele counts (copy numbers >1) which have more overlapping reads proportional to the number of alleles and to the sequenced depth. The specific steps for FOD calculation are detailed as follows:

1. **Read pair filtering and retrieval** For a given coordinate range (e.g., chr1:20000-30000), "samtools view" is used with "-F 3996 -q 30" to retrieve high-quality read pairs within the region from the single-cell bam. From the filtered reads, we retain the following attributes for the next step: start position of the first read in pair, fragment length, Compact Idiosyncratic Gapped Alignment Report (CIGAR) string, and read ID.
2. **Fragment coordinate adjustment** We adjust all fragment end coordinates by subtracting 10 base pairs to align with the staggered cut patterns often produced by Tn5-transposase, which are typically 9 or 10 base pairs in length [2].
3. **Filtering** We implemented the following filters to remove false-positive overlapping cases due to poor mapping quality and sequencing artifacts. First, we discard fragments whose CIGAR string indicates soft clipping ("S"), deletion ("D"), or insertion ("I") events, as these may represent misalignment errors. Second, we remove fragments with identical start or end positions which could be indicative of technical artifacts that arise in PCR or repetitive genomic regions, rather than distinct molecular identities.

4. **Efficient identification of overlapping fragments using binary search** The most naïve approach to search for overlapping fragments would be a double iteration over the read pairs (or fragments), where for each fragment, its location is compared with every other fragment in the region of interest to check for intersection. Overlap occurs if the start of one fragment is within another fragment, or vice versa. Specifically, for two fragments A and B, an overlap exists if $A_{\text{start}} \leq B_{\text{end}}$, and $A_{\text{end}} \geq B_{\text{start}}$. Since the naïve method requires two nested loops, the time complexity is $O(n^2)$ which can be prohibitively expensive as the number of fragments increases when the query region is large or was sequenced to high depth. To improve efficiency, we implemented a sorting and binary search-based model. First, we sort the start and end (i.e., left and right) coordinates of the fragments within the region of interest. After sorting, we perform binary search using the python "bisect" package [3] to find the positions where a fragment's start or end would be inserted into the sorted list of ends or starts. Specifically, we find how many fragments have their right end less than the current fragment's left start, and how many have their left start greater than the current fragment's right end. The number of overlaps for a given fragment is then the total number of fragments minus those on the left and right that do not overlap with it, and minus itself. This procedure takes a time complexity of $O(nlogn)$, making it significantly more efficient than the naïve approach.
5. **Calculation of FOD** The FOD is then computed as : $FOD = \frac{N_{\text{overlap}}}{(N_{\text{total}} - N_{\text{overlap}})}$,
where $N_{\text{overlap}}$ represents the number of unique overlapping instances and $N_{\text{total}}$ is the total number of reads in the region. This calculation yields a ratio that increases proportionally with copy number, as higher copy numbers provide more opportunities for fragment overlap. In cases where the denominator is zero, which tends to happen when the total number of reads per cell falls below 10M reads, the FOD is recorded as undefined, and the call cannot be validated.

#### Performance assessment of FOD using synthetic diploid X construct

We evaluated the performance of FOD using a synthetic diploid chromosome X benchmarking dataset derived from male samples. Six read depth categories, spanning 10–150M reads, were defined on a log10 scale ([7–8.000, 8.000–8.125, 8.125–8.250, 8.250–8.375, 8.375–8.500, >8.500]). In each depth category, three single cell genomes from a male subject UMB5871 with no detected CNVs (by HiScanner) were selected randomly. These cells' read depth ratio tracks were visually inspected to ensure the X chromosomes were CNV-free. Read records for chromosome X from these cells in each depth category were retrieved and reads in PARs or those failing quality thresholds (e.g., fragment length <50 or >450 bp) were excluded. Haploid (CN1) states were represented by reads from one cell, while diploid (CN2) and triploid (CN3) states were generated by concatenating reads from two and three cells, respectively. Simulated CNV regions were generated by randomly selecting 100 non-overlapping regions for 12 predefined CNV lengths (ranging from 2 Mb to 24 Mb) across the entire span of the X chromosome, excluding PARs.

FOD Z-scores were calculated for predefined CNV regions using diploid mean and standard deviation values derived from diploid segments (i.e., regions with predicted copy number equal to two). Simulated CNVs were divided into loss (CN1 vs. CN2) and gain (CN3 vs. CN2) datasets. Binary labels were assigned to segments based on their true CNV state: CN1 and CN3 were labeled as 1 (CNV present), and CN2 as 0 (CNV absent). Predictions were made using Z-score thresholds: segments with Z-scores $<$ -1.96 (loss) or $>$ 1.96 (gain) were classified as CNVs. Precision, or positive predictive value (PPV), was calculated as: $PPV = \frac{TP}{TP+FP}$ where true positives (TP) are segments correctly identified as CNVs (e.g., Z-scores $<$ -1.96 for CN1 or $>$ 1.96 for CN3), and false positives (FP) are segments incorrectly classified as CNVs (e.g., Z-scores meeting thresholds for regions that are actually CN2). If $TP + FP = 0$, PPV was set to 0. PPV was computed separately for each read depth category and CNV length to evaluate performance across varying genomic contexts.

#### Apoptosis assay for neuronal progenitor cell control

NPCs were developed using Human WT iPSC line "280" derived from healthy male fibroblasts generated by Boston Children's Hospital hESC Core [4]. These iPSC were differentiated into NPCs using STEMdiff SMADi Neural Induction Kit (Cat: 08582 and "Monolayer Culture Protocol"). Apoptosis was induced in the cell lines using staurosporin 1$\mu$M for 12 hours. Cells were stained with Annexin V (BioRad Annexin V Kit ANNEX200APC) and Propidium Iodide (ThermoFisher P1304MP) according to manufacturer instructions. Flow cytometry was performed to identify two populations of single nuclei: 1) no nuclear disruption, characterized by low propidium iodide positivity and high Annexin V positivity, and 2) with nuclear disruption, represented by high propidium iodide fluorescence positivity and high Annexin V fluorescence. Individual nuclei were sorted into microwells, and scWGS and CNV calling were subsequently performed using the same methods as for human tissue.

#### Single-cell RNA sequencing and analysis for neuronal proportion estimate in cortical plates

Isolation of neuronal nuclei was performed as described above. Nuclei were sorted directly into buffer for 10X Genomics Single Cell 3' v3.1 kit (PN-1000128). GEMs were generated per manufacturer's instructions. Double sided size selection using SPRIselect was performed to optimize library size. Sequencing data analysis was performed using Seurat (version 5.0.1) [5] and Signac (version 1.12.0) [6] R packages. Cells with count <1000 or >3 median absolute deviations (MADs) from the median were removed to exclude outliers. Cell type annotation was performed by mapping each sample to a reference scRNAseq dataset. We performed cell type label transfer using the Symphony R library (v0.1.0) for the remaining non-reference "query" samples. First, we used Symphony's buildReference function to create a reference from our selected reference samples. Then, we used the mapQuery function to integrate the new query data into the reference samples, projecting the query samples onto the reference UMAPs. Finally, Symphony's knnPredict function was applied to predict the cell type annotations of query cells from the reference using the k-nearest neighbors (kNN) method. This approach enabled comprehensive cell type annotation for all samples in the multiome dataset.

#### Isolation and processing of micronuclei through serial sucrose gradient fractionation

Tissues were homogenized in 5 ml pre-chilled lysis buffer (10mM Tris-HCl, 2mM Mg-acetate, 3 mM CaCl2, 0.32M sucrose, 0.1mM EDTA, 1mM DTT, 0.1% NP-40, 0.15mM spermine, 0.75mM spermidine and 10 $\mu$g/ml cytochalasin B, pH 8.5). The homogenate was subsequently filtered through a 40uM filter, mixed with 5mL 1.8M sucrose buffer (10mM Tris HCl pH 8.5, 5mM MgAc2, 0.1mM EDTA, 1mM DTT, 0.3% BSA, 0.15mM spermine, 0.75mM spermidine, 1.8M sucrose). The mixture was then layered on top of a sucrose buffer (20ml 1.8M sucrose (bottom), 15ml 1.6M sucrose (middle)) and centrifuged in Thermo Scientific™ Sorvall LYNX 6000 Superspeed Centrifuge (Catalog No.75-006-590) at 2,400 rpm at 4°C. Subsequently, five 3ml fractions were collected sequentially starting from the top layer. Each fraction was stained with DAPI and visually checked under a microscope to identify the two fractions enriched in micronuclei. Both fractions were then layered onto 4.7ml of 1.8M sucrose buffer and centrifuged at max speed for 90 minutes at 4°C. After discarding the supernatant, the pellet was resuspended in 1ml 1xPBS, layered on top of a linear sucrose gradient (1.0 to 1.8M sucrose buffer), and centrifuged at 1,800 rpm for 15 minutes at 4°C. Then, 1ml fractions were collected sequentially from the top. Four micronuclei-enriched fractions were combined, diluted 5-fold in 1x PBS, and centrifuged at 2,400 rpm for 15 minutes at 4°C. About 500$\mu$l supernatant was kept to avoid disturbing the micronuclei pellet. The resuspension underwent phenol-chloroform extraction, and 1ul of the resulting DNA suspension was input for low-input library Nextera-based library amplification. We note that we utilize our at home loaded Tn5 transposase for library preparation, i.e. Step 2 of TbA library preparation, as per "Single-cell whole genome library preparation" (Methods).

#### Informatic analysis of paired micronuclei and nuclei genome sequencing

Sequenced micronuclear and nuclear libraries were analyzed to determine genomic regions that may be over- or under-enriched in the micronuclei fraction compared to the nuclear fraction. BIC-seq2 [7] was used to quantify read counts and identify breakpoint candidates. This analysis was conducted using a mappable bin size of 50kb. Subsequently, the MN/N log ratio was computed using the logarithm base 2 of the ratio of micronuclei (MN) read counts over nuclei (N) read counts in each chromosome for a given sample.

We estimated the change in the ratio of micronucleus-to-nucleus read depth over time using a generalized linear model (GLM) with age (post-conception weeks) and chromosomal GC content (z-scored) as predictors of $|\log_2(\mathrm{MN/N})|$, treated as an exponentially distributed random variable following quantile–quantile analysis. We used the `glm` function in R with a Gamma family and log link. To confirm model selection, we compared three nested models—a null model, a model including age only, and a model including both age and chromosomal GC content—using AIC and deviance as goodness-of-fit criteria. The model including both predictors provided the best fit, minimizing AIC ($-495$ vs. $-474$ vs. $-455$) and deviance (215 vs. 237 vs. 261).

#### Informatic meta-analysis of age-related CNV trends across technologies

To evaluate the association between age and the proportion of cells harboring copy number variants in our study and prior publications (Table S2), we used a linear mixed-effects model implemented in Python using the `mixedlm` function from the `statsmodels` package (v0.14). The model was specified as:

$$\texttt{pct\_cnv} \sim \texttt{age} + \texttt{cell\_count} + (1 \mid \texttt{study}),$$

where `pct_cnv` is the percentage of cells carrying any CNV, `age` is donor age in years, and `cell_count` is the mean-centered number of cells sequenced per individual. `study` was included as a random intercept to account for systematic differences in baseline CNV detection rates arising from variation in sequencing platform, CNV calling pipeline, and tissue processing across cohorts. Model parameters were estimated using restricted maximum likelihood. TbA and PTA data from this study were treated as separate studies in this grouping.

#### Immunohistochemistry

For frozen postmortem brain tissue: 10$\mu$m cryosections of brain tissue were placed on Superfrost slides (Fisher) for immunohistochemistry. The sample was first fixed in 4% PFA for 15 minutes at room temperature, followed by permeabilization with 0.5% Triton-X in PBS for 1 hour and blocked with 10% donkey serum in PBS and 0.05% Triton-X (PBST) for 30 minutes. Primary mouse anti-SATB2 (Abcam, ab9244) diluted 1:500 in blocking solution was applied to the sections overnight at 4°C. After 5 washes with PBST at RT for 5 minutes, secondary antibody (Donkey anti-Mouse Alexa Fluor 488) and DAPI diluted in blocking solution were applied to the sections overnight at 4°C. Finally, sections were washed with PBST for a minimum of 5 times before mounting with Vectashield Vibrance Antifade Mounting Medium. Images were captured at 63X magnification by a Zeiss LSM 980 confocal microscope. Z-stack function was used to image a 10 $\mu$m thickness. Sample images were prepared in ImageJ software.

For fractions obtained from serial sucrose gradient fractionation: 200 µL from each fraction were fixed in 2% paraformaldehyde (PFA) for 10 min at room temperature. Samples were pelleted by centrifugation at 500 $\times$ g and washed twice with 1$\times$ phosphate-buffered saline (PBS). Pellets were resuspended in 100 µL PBS then the suspension was deposited onto Superfrost slides (Fisher) and allowed to air-dry overnight at room temperature. Slides were permeabilized and blocked in blocking buffer containing 0.3% Triton X-100 and 0.75% normal goat serum (Abcam, ab7481) diluted in PBS for 90 min at room temperature. Primary antibody incubation was performed overnight at

4 °C in a humidified chamber using rabbit anti–Lamin A/Lamin B1/Lamin C antibody (Abcam, ab108922), diluted 1:200 in blocking buffer. Following three washes in PBS (5 min each, RT), slides were incubated with goat anti-rabbit Alexa Fluor 555 secondary antibody (1:250 dilution in blocking buffer) for 1 h at room temperature in the dark. Slides were washed three times in PBS (7 min each, room temperature) and mounted using Fluoromount-G® mounting medium containing DAPI (SouthernBiotech). Images were acquired using a Zeiss LSM 980 confocal microscope equipped with a 63× oil-immersion objective.

#### Micronuclei enumeration

Z-stack nuclear images were imported into the MATLAB-based program CAMDi [8] for the quantification of micronuclei. The nuclear images were color-coded in blue and red, with an overlap area threshold set at 0.5. The imported data underwent binarization, with a threshold established at 25.0. Following the creation of binary images, the merged areas were extracted based on this threshold, enabling the quantification of both the number and volume of micronuclei. Micronuclei were defined as having a diameter greater than 0.1 um and less than 1.0 um. For unbiased counting of main nuclei in fluorescence images stained with DAPI, we employed Cellpose, a deep learning-based segmentation algorithm (version 2.2.2, CPx model, cell diameter = 30 pixels). Additionally, for manual quantification, regions of interest (ROIs) for main nuclei and micronuclei were delineated and determined using the ROI Manager in FIJI ImageJ (version 2.14.0/1.54f), where nuclei with a longest diameter of 1 um or less were classified as micronuclei.

#### Generation of word cloud

Gene Ontology Cellular Component (GO-CC) terms that were significantly downregulated in aneuploid excitatory neurons from postnatal samples (Table S1; Bonferroni-corrected p-value $< 0.05$) were used to generate a word cloud visualization. Terms were first filtered to retain only those with absolute observed differences above the 80th percentile and negative observed differences, representing strongly down-regulated pathways. Gene set names were processed by removing the "GO_CC" prefix and replacing underscores with spaces. The processed terms were then used to generate a word cloud using the Free Word Cloud Generator [9] where the size of each term reflects its frequency among the significant gene sets.

### Supplemental Tables

#### **Table S1** Summary of samples.

Sample table of all prenatal, postnatal, and cell line TbA and PTA libraries used in this study. NPC: neural progenitor cell line. *Low rate includes exclusion of replicating NPC. gw, gestational weeks. y, years.

#### **Table S2.** CNV Table

Genomic coordinates of CNV breakpoints for all neurons and donors in this study.

#### **Table S3.** Postnatal neuron CNV meta-table

Consolidated results of single neuron CNV detection in postnatal neurons across different studies and scWGA technologies.

**Table S4.** Micronuclei enumeration.

Micronuclei and primary nuclei enumeration in the cortical plate. Enumerated across 5 different images.

**Table S5.** Micronuclei sequencing.

Sequencing statistics for paired micronuclei (MN) and nuclei (N) isolated by sucrose gradient fractionation.

**Table S6.** Differential gene module scores.

Only gene modules with corrected p-value $< 0.05$ are shown. Gene modules with absolute observed difference $> 0.5$ are highlighted in yellow.

**Supplementary Table 1: Sample table of all prenatal, postnatal, and cell line TbA and PTA libraries.**

| Donor ID | Age group | Library type | Sex | Age | Sequenced cell number | Leiden 0 (%) | Final QC pass rate (%) | Final cell number | PMI |
|---|---|---|---|---|---|---|---|---|---|
| UMB4827 | Prenatal | PTA | Male | 14gw | 88 | 80 | 80 | 70 | 5 |
| UMB1117 | Prenatal | PTA | Male | 15gw | 67 | 84 | 84 | 56 | 2 |
| UMB1367 | Prenatal | PTA | Male | 15gw | 74 | 70 | 70 | 52 | 1 |
| UMB1932 | Prenatal | PTA | Male | 21gw | 73 | 74 | 74 | 54 | 24 |
| H29648 | Prenatal | PTA | Female | 18gw | 76 | 76 | 76 | 58 | |
| UMB4388 | Postnatal | PTA | Female | 0.4y | 74 | 96 | 96 | 71 | 53 |
| UMB1284 | Postnatal | PTA | Female | 3y | 42 | 60 | 60 | 25 | 11 |
| UMB6032 | Postnatal | PTA | Male | 4y | 80 | 64 | 64 | 51 | 25 |
| UMB1465 | Postnatal | PTA | Male | 17.5y | 58 | 69 | 69 | 40 | 4 |
| UMB936 | Postnatal | PTA | Female | 49y | 54 | 83 | 83 | 45 | 7 |
| UMB5451 | Postnatal | PTA | Male | 57y | 72 | 92 | 92 | 66 | 18 |
| UMB5828 | Postnatal | PTA | Female | 66y | 73 | 97 | 97 | 71 | 25 |
| UMB4827 | Prenatal | TbA | Male | 14gw | 93 | 98 | 85 | 79 | 5 |
| UMB1117 | Prenatal | TbA | Male | 15gw | 85 | 84 | 38 | 32 | 2 |
| UMB1367 | Prenatal | TbA | Male | 15gw | 81 | 85 | 64 | 52 | 1 |
| UMB14 | Prenatal | TbA | Female | 18gw | 40 | 98 | 90 | 36 | 3 |
| UMB219 | Prenatal | TbA | Female | 18gw | 95 | 80 | 56 | 53 | 1 |
| H29648 | Prenatal | TbA | Female | 18gw | 86 | 98 | 92 | 79 | |
| UMB1031 | Prenatal | TbA | Female | 20gw | 64 | 88 | 78 | 50 | 2 |
| UMB1932 | Prenatal | TbA | Male | 21gw | 92 | 86 | 39 | 36 | 24 |
| UMB4417 | Prenatal | TbA | Male | 28gw | 77 | 91 | 69 | 53 | 45 |
| UMB4428 | Postnatal | TbA | Male | 0.2y | 81 | 99 | 90 | 73 | |
| UMB1864 | Postnatal | TbA | Female | 2y | 78 | 87 | 76 | 59 | 8 |
| UMB5871 | Postnatal | TbA | Male | 2y | 91 | 100 | 100 | 91 | 12 |
| UMB4782 | Postnatal | TbA | Male | 18y | 86 | 100 | 85 | 73 | 17 |
| UMB5451 | Postnatal | TbA | Male | 57y | 76 | 78 | 49 | 37 | 18 |
| E853 | Postnatal | TbA | Female | 82y | 123 | 82 | 60 | 74 | 13 |
| MAYO047 | Postnatal | TbA | Female | 90y | 79 | 96 | 90 | 71 | 13 |
| Disrupted nuclei | NPC | TbA | - | - | 54 | 59* | - | 32 | - |
| Intact nuclei | NPC | TbA | - | - | 71 | 66* | - | 47 | - |

Supplementary Table 2: CNV Table. Available upon request due to file size limits for BioRxiv.

**Supplementary Table 3: Summary of single-cell CNV detection in postnatal neurons across different stu**

| Study | Platform | Age (years) | % Neurons with CNV | Cell Count |
|---|---|---|---|---|
| This study | TbA | 0.2 | 42.46575342 | 73 |
| This study | PTA | 0.4 | 15.49295775 | 71 |
| This study | TbA | 2 | 20.87912088 | 91 |
| This study | TbA | 2 | 49.15254237 | 59 |
| This study | PTA | 3.3 | 12 | 25 |
| This study | PTA | 4 | 45.09803922 | 51 |
| This study | PTA | 17 | 20 | 40 |
| This study | TbA | 18 | 41.09589041 | 73 |
| This study | PTA | 49 | 13.63636364 | 44 |
| This study | TbA | 57 | 16.21621622 | 37 |
| This study | PTA | 57 | 37.87878788 | 66 |
| This study | PTA | 66 | 16.90140845 | 71 |
| This study | TbA | 82 | 16.21621622 | 74 |
| This study | TbA | 90 | 18.30985915 | 71 |
| Cai 2014 | GenomePlex | 42 | 36.8 | 19 |
| Chronister 2019 | PicoPLEX | 0.36 | 30.4 | 46 |
| Chronister 2019 | PicoPLEX | 26 | 23.1 | 108 |
| Chronister 2019 | PicoPLEX | 49 | 11.1 | 99 |
| Chronister 2019 | PicoPLEX | 49 | 7.7 | 26 |
| Chronister 2019 | PicoPLEX | 86 | 4 | 101 |
| Chronister 2019 | PicoPLEX | 86 | 8.7 | 46 |
| Chronister 2019 | PicoPLEX | 95 | 6.7 | 120 |
| Knouse 2014 | GenomePlex | 48 | 0 | 21 |
| Knouse 2014 | GenomePlex | 52 | 18.2 | 22 |
| Knouse 2014 | GenomePlex | 70 | 20 | 20 |
| Knouse 2014 | GenomePlex | 68 | 13 | 23 |
| McConnell 2013 | GenomePlex | 20 | 40 | 50 |
| McConnell 2013 | GenomePlex | 24 | 38.9 | 18 |
| McConnell 2013 | GenomePlex | 26 | 22 | 41 |
| Sun 2024 | 10X Genomics | 49 | 10.8 | 2097 |
| van den Bos 2016 | Strand-seq | 69 | 3.8 | 78 |
| van den Bos 2016 | Strand-seq | 74 | 8.6 | 70 |
| van den Bos 2016 | Strand-seq | 82 | 9.3 | 43 |
| Zeliha Gšzde Turan 2022 | GenomePlex | 78 | 0 | 56 |
| Zeliha Gšzde Turan 2022 | GenomePlex | 69 | 0 | 127 |
| Zeliha Gšzde Turan 2022 | GenomePlex | 60 | 0 | 61 |
| Zeliha Gšzde Turan 2022 | GenomePlex | 92 | 1.61 | 62 |
| Zeliha Gšzde Turan 2022 | GenomePlex | 68 | 0 | 63 |
| Zeliha Gšzde Turan 2022 | GenomePlex | 52 | 0.7 | 149 |
| Zeliha Gšzde Turan 2022 | GenomePlex | 82 | 2.1 | 94 |

**Supplementary Table 4: Micronuclei and primary nuclei enumeration in the cortical plat**

| | Micronuclei (manual) | Main nuclei (Cellpose) | Main nuclei (manual) |
|---|---|---|---|
| Image 1 | 26 | 265 | 251 |
| Image 2 | 24 | 374 | 324 |
| Image 3 | 15 | 457 | 361 |
| Image 4 | 11 | 465 | 416 |
| Image 5 | 27 | 443 | 349 |
| **Average** | **20.6** | **400.8** | **340.2** |

**Supplementary Table 5: Sequencing statistics for paired micronuclei (MN) and nuclei (N) isolated by s[illegible]**

| Donor ID | MN Mean Depth | N Mean Depth | MN Mean Coverage (%) | N Mean Coverage (%) |
|---|---|---|---|---|
| UMB1932 | 1.85 | 2.01 | 67.25% | 70.40% |
| UMB4428 | 1.58 | 3.19 | 62.69% | 80.88% |
| UMB4942 | 1.43 | 2.16 | 47.67% | 62.16% |
| UMB5621 | 0.93 | 2.97 | 34.95% | 69.42% |
| UMB5926 | 1.36 | 2.17 | 46.02% | 63.44% |
| UMB5871 | 0.43 | 2.04 | 17.24% | 59.65% |
| UMB936 | 2.75 | 3.05 | 78.62% | 80.01% |
| UMB6032 | 0.07 | 0.71 | 5.78% | 33.03% |

**Supplementary Table 6: Differential gene module scores across all samples. Only gene modules with corrected p-value < 0.05 are shown. Gene modules with absolute observed differ**

| Gene_Set | Age group | Pathway database | Cell type | Observed Difference | Absolute Observed Difference | P Value | P value corrected |
|---|---|---|---|---|---|---|---|
| GOCC_CHROMOSOME_CENTROMERIC_REGION | fetal | c5_gocc | Progenitor | 0.0781 | 0.0781 | 0 | 0 |
| GOCC_CONDENSED_CHROMOSOME_CENTROM | fetal | c5_gocc | Progenitor | 0.0760 | 0.0760 | 0 | 0 |
| GOCC_CONDENSED_CHROMOSOME1 | fetal | c5_gocc | Progenitor | 0.0760 | 0.0760 | 0 | 0 |
| GOCC_SPINDLE1 | fetal | c5_gocc | Progenitor | 0.0558 | 0.0558 | 0 | 0 |
| GOCC_CUL2_RING_UBIQUITIN_LIGASE_COMPL | fetal | c5_gocc | ExN | -0.0233 | 0.0233 | 0 | 0 |
| WU_APOPTOSIS_BY_CDKN1A_VIA_TP531 | fetal | keywords_c2 | Progenitor | 0.1738 | 0.1738 | 0.0001 | 0.0150 |
| TANG_SENESCENCE_TP53_TARGETS_DN1 | fetal | keywords_c2 | Progenitor | 0.1679 | 0.1679 | 0 | 0 |
| REACTOME_DNA_DAMAGE_TELOMERE_STRES | fetal | keywords_c2 | Progenitor | 0.0686 | 0.0686 | 0.0003 | 0.0450 |
| HOLLMANN_APOPTOSIS_VIA_CD40_DN1 | fetal | keywords_c2 | InN | -0.0077 | 0.0077 | 0.0002 | 0.0300 |
| GOCC_POSTSYNAPTIC_DENSITY_MEMBRANE1 | postnatal | c5_gocc | Non-neuron | -0.1967 | 0.1967 | 0 | 0 |
| GOCC_NEURON_TO_NEURON_SYNAPSE1 | postnatal | c5_gocc | Non-neuron | -0.1346 | 0.1346 | 0 | 0 |
| GOCC_SYNAPTIC_MEMBRANE1 | postnatal | c5_gocc | Non-neuron | -0.1320 | 0.1320 | 0 | 0 |
| GOCC_POSTSYNAPSE1 | postnatal | c5_gocc | Non-neuron | -0.1025 | 0.1025 | 0 | 0 |
| GOCC_NEURON_PROJECTION1 | postnatal | c5_gocc | Non-neuron | -0.0769 | 0.0769 | 0 | 0 |
| GOCC_HISTONE_METHYLTRANSFERASE_COMP | postnatal | c5_gocc | Non-neuron | 0.0368 | 0.0368 | 0 | 0 |
| GOCC_MLL1_2_COMPLEX1 | postnatal | c5_gocc | Non-neuron | 0.0223 | 0.0223 | 0 | 0 |
| GOCC_NUCLEAR_STRESS_GRANULE1 | postnatal | c5_gocc | ExN | -4.3666 | 4.3666 | 0 | 0 |
| GOCC_JUXTAPARANODE_REGION_OF_AXON1 | postnatal | c5_gocc | ExN | -3.2932 | 3.2932 | 0 | 0 |
| GOCC_AXOLEMMA1 | postnatal | c5_gocc | ExN | -2.2408 | 2.2408 | 0 | 0 |
| GOCC_CYTOSKELETON_OF_PRESYNAPTIC_AC | postnatal | c5_gocc | ExN | -2.0701 | 2.0701 | 0 | 0 |
| GOCC_PRESYNAPTIC_CYTOSKELETON1 | postnatal | c5_gocc | ExN | -1.5181 | 1.5181 | 0 | 0 |
| GOCC_ASYMMETRIC_GLUTAMATERGIC_EXCITA | postnatal | c5_gocc | ExN | -1.4902 | 1.4902 | 0 | 0 |
| GOCC_POTASSIUM_CHANNEL_COMPLEX1 | postnatal | c5_gocc | ExN | -1.4425 | 1.4425 | 0 | 0 |
| GOCC_MAIN_AXON1 | postnatal | c5_gocc | ExN | -1.3269 | 1.3269 | 0 | 0 |
| GOCC_AXON_INITIAL_SEGMENT1 | postnatal | c5_gocc | ExN | -1.2910 | 1.2910 | 0 | 0 |
| GOCC_SYNAPTIC_CLEFT1 | postnatal | c5_gocc | ExN | -1.2011 | 1.2011 | 0 | 0 |
| GOCC_POSTSYNAPTIC_SPECIALIZATION_MEMI | postnatal | c5_gocc | ExN | -1.1162 | 1.1162 | 0 | 0 |
| GOCC_SYNAPTIC_MEMBRANE1 | postnatal | c5_gocc | ExN | -1.1109 | 1.1109 | 0 | 0 |
| GOCC_POSTSYNAPTIC_DENSITY_MEMBRANE1 | postnatal | c5_gocc | ExN | -1.1038 | 1.1038 | 0 | 0 |
| GOCC_HIPPOCAMPAL_MOSSY_FIBER_TO_CA3_ | postnatal | c5_gocc | ExN | -1.0586 | 1.0586 | 0 | 0 |
| GOCC_POSTSYNAPTIC_MEMBRANE1 | postnatal | c5_gocc | ExN | -0.9790 | 0.9790 | 0 | 0 |
| GOCC_CATION_CHANNEL_COMPLEX1 | postnatal | c5_gocc | ExN | -0.9581 | 0.9581 | 0 | 0 |
| GOCC_PRESYNAPTIC_MEMBRANE1 | postnatal | c5_gocc | ExN | -0.9402 | 0.9402 | 0 | 0 |
| GOCC_EXCITATORY_SYNAPSE1 | postnatal | c5_gocc | ExN | -0.9208 | 0.9208 | 0 | 0 |
| GOCC_NEURON_TO_NEURON_SYNAPSE1 | postnatal | c5_gocc | ExN | -0.8769 | 0.8769 | 0 | 0 |
| GOCC_PRESYNAPTIC_ACTIVE_ZONE_CYTOPLA | postnatal | c5_gocc | ExN | -0.8652 | 0.8652 | 0 | 0 |
| GOCC_COSTAMERE1 | postnatal | c5_gocc | ExN | -0.8417 | 0.8417 | 0 | 0 |
| GOCC_CKM_COMPLEX1 | postnatal | c5_gocc | ExN | 0.8362 | 0.8362 | 0 | 0 |
| GOCC_POSTSYNAPTIC_SPECIALIZATION1 | postnatal | c5_gocc | ExN | -0.8281 | 0.8281 | 0 | 0 |
| GOCC_POSTSYNAPSE1 | postnatal | c5_gocc | ExN | -0.7509 | 0.7509 | 0 | 0 |
| GOCC_MONOATOMIC_ION_CHANNEL_COMPLEX | postnatal | c5_gocc | ExN | -0.7297 | 0.7297 | 0 | 0 |
| GOCC_GLUTAMATERGIC_SYNAPSE1 | postnatal | c5_gocc | ExN | -0.7136 | 0.7136 | 0 | 0 |
| GOCC_NEURON_SPINE1 | postnatal | c5_gocc | ExN | -0.6626 | 0.6626 | 0 | 0 |
| GOCC_NEURON_PROJECTION_MEMBRANE1 | postnatal | c5_gocc | ExN | -0.6391 | 0.6391 | 0 | 0 |
| GOCC_X_CHROMOSOME1 | postnatal | c5_gocc | ExN | 0.6281 | 0.6281 | 0 | 0 |
| GOCC_PARASPECKLES1 | postnatal | c5_gocc | ExN | 0.6087 | 0.6087 | 0 | 0 |
| GOCC_DENDRITIC_TREE1 | postnatal | c5_gocc | ExN | -0.5987 | 0.5987 | 0 | 0 |
| GOCC_SYNAPSE1 | postnatal | c5_gocc | ExN | -0.5869 | 0.5869 | 0 | 0 |
| GOCC_AXON1 | postnatal | c5_gocc | ExN | -0.5869 | 0.5869 | 0 | 0 |
| GOCC_PRESYNAPTIC_ACTIVE_ZONE1 | postnatal | c5_gocc | ExN | -0.5729 | 0.5729 | 0 | 0 |
| GOCC_CATENIN_COMPLEX1 | postnatal | c5_gocc | ExN | -0.5683 | 0.5683 | 0 | 0 |
| GOCC_SOMATODENDRITIC_COMPARTMENT1 | postnatal | c5_gocc | ExN | -0.5669 | 0.5669 | 0 | 0 |
| GOCC_CLATHRIN_COMPLEX1 | postnatal | c5_gocc | ExN | 0.5627 | 0.5627 | 0 | 0 |
| GOCC_STEREOCILIA_ANKLE_LINK_COMPLEX1 | postnatal | c5_gocc | ExN | 0.5545 | 0.5545 | 0 | 0 |
| GOCC_SPECTRIN_ASSOCIATED_CYTOSKELETO | postnatal | c5_gocc | ExN | -0.5541 | 0.5541 | 0 | 0 |
| GOCC_TRANSPORTER_COMPLEX1 | postnatal | c5_gocc | ExN | -0.5529 | 0.5529 | 0 | 0 |
| GOCC_NEURON_PROJECTION1 | postnatal | c5_gocc | ExN | -0.5432 | 0.5432 | 0 | 0 |
| GOCC_NEUROTRANSMITTER_RECEPTOR_COMI | postnatal | c5_gocc | ExN | -0.5373 | 0.5373 | 0 | 0 |
| GOCC_AXON_HILLOCK1 | postnatal | c5_gocc | ExN | -0.5266 | 0.5266 | 0 | 0 |
| GOCC_VOLTAGE_GATED_CALCIUM_CHANNEL_ | postnatal | c5_gocc | ExN | -0.5249 | 0.5249 | 0 | 0 |
| GOCC_CELL_BODY1 | postnatal | c5_gocc | ExN | -0.4880 | 0.4880 | 0 | 0 |
| GOCC_PRESYNAPSE1 | postnatal | c5_gocc | ExN | -0.4750 | 0.4750 | 0 | 0 |
| GOCC_NUCLEAR_CYCLIN_DEPENDENT_PROTE | postnatal | c5_gocc | ExN | 0.4690 | 0.4690 | 0 | 0 |
| GOCC_CELL_CORTEX_REGION1 | postnatal | c5_gocc | ExN | -0.4616 | 0.4616 | 0 | 0 |
| GOCC_PERIKARYON1 | postnatal | c5_gocc | ExN | -0.4608 | 0.4608 | 0 | 0 |
| GOCC_STEREOCILIUM_MEMBRANE1 | postnatal | c5_gocc | ExN | 0.4434 | 0.4434 | 0 | 0 |
| GOCC_SYNAPSE_ASSOCIATED_EXTRACELLUL | postnatal | c5_gocc | ExN | 0.4409 | 0.4409 | 0 | 0 |
| GOCC_NEUROMUSCULAR_JUNCTION1 | postnatal | c5_gocc | ExN | -0.4375 | 0.4375 | 0 | 0 |
| GOCC_PROXIMAL_DENDRITE1 | postnatal | c5_gocc | ExN | -0.4341 | 0.4341 | 0 | 0 |
| GOCC_PLASMA_MEMBRANE_REGION1 | postnatal | c5_gocc | ExN | -0.4290 | 0.4290 | 0 | 0 |
| GOCC_T_TUBULE1 | postnatal | c5_gocc | ExN | -0.4219 | 0.4219 | 0 | 0 |
| GOCC_MITOCHONDRIAL_CRISTA_JUNCTION1 | postnatal | c5_gocc | ExN | 0.4204 | 0.4204 | 0 | 0 |
| GOCC_ADHERENS_JUNCTION1 | postnatal | c5_gocc | ExN | -0.4179 | 0.4179 | 0 | 0 |
| GOCC_TORC2_COMPLEX1 | postnatal | c5_gocc | ExN | 0.4170 | 0.4170 | 0 | 0 |
| GOCC_AMPA_GLUTAMATE_RECEPTOR_COMPL | postnatal | c5_gocc | ExN | -0.4145 | 0.4145 | 0 | 0 |
| GOCC_SCHAFFER_COLLATERAL_CA1_SYNAPS | postnatal | c5_gocc | ExN | -0.4084 | 0.4084 | 0 | 0 |
| GOCC_PARALLEL_FIBER_TO_PURKINJE_CELL_ | postnatal | c5_gocc | ExN | -0.4067 | 0.4067 | 0 | 0 |
| GOCC_CALCIUM_CHANNEL_COMPLEX1 | postnatal | c5_gocc | ExN | -0.4055 | 0.4055 | 0 | 0 |
| GOCC_COLLAGEN_TYPE_IV_TRIMER1 | postnatal | c5_gocc | ExN | 0.4045 | 0.4045 | 0 | 0 |
| GOCC_BARR_BODY1 | postnatal | c5_gocc | ExN | 0.4035 | 0.4035 | 0 | 0 |
| GOCC_CENTRIOLAR_SUBDISTAL_APPENDAGE1 | postnatal | c5_gocc | ExN | 0.3971 | 0.3971 | 0 | 0 |
| GOCC_CELL_CELL_JUNCTION1 | postnatal | c5_gocc | ExN | -0.3902 | 0.3902 | 0 | 0 |
| GOCC_PLASMA_MEMBRANE_PROTEIN_COMPLI | postnatal | c5_gocc | ExN | -0.3883 | 0.3883 | 0 | 0 |
| GOCC_COMPACT_MYELIN1 | postnatal | c5_gocc | ExN | 0.3828 | 0.3828 | 0 | 0 |
| GOCC_SARCOLEMMA1 | postnatal | c5_gocc | ExN | -0.3693 | 0.3693 | 0 | 0 |
| GOCC_DENDRITIC_SHAFT1 | postnatal | c5_gocc | ExN | -0.3667 | 0.3667 | 0 | 0 |

| | | | | | | | |
|---|---|---|---|---|---|---|---|
| GOCC_POLAR_MICROTUBULE1 | postnatal | c5_gocc | ExN | 0.3658 | 0.3658 | 0 | 0 |
| GOCC_NSL_COMPLEX1 | postnatal | c5_gocc | ExN | 0.3646 | 0.3646 | 0 | 0 |
| GOCC_NEURON_PROJECTION_CYTOPLASM1 | postnatal | c5_gocc | ExN | -0.3617 | 0.3617 | 0 | 0 |
| GOCC_STEREOCILIA_COUPLING_LINK1 | postnatal | c5_gocc | ExN | 0.3499 | 0.3499 | 0 | 0 |
| GOCC_AXON_CYTOPLASM1 | postnatal | c5_gocc | ExN | -0.3487 | 0.3487 | 0 | 0 |
| GOCC_FLEMMING_BODY1 | postnatal | c5_gocc | ExN | -0.3404 | 0.3404 | 0 | 0 |
| GOCC_MIGRASOME1 | postnatal | c5_gocc | ExN | 0.3359 | 0.3359 | 0 | 0 |
| GOCC_PERICENTRIOLAR_MATERIAL1 | postnatal | c5_gocc | ExN | 0.3334 | 0.3334 | 0 | 0 |
| GOCC_TOR_COMPLEX1 | postnatal | c5_gocc | ExN | 0.3300 | 0.3300 | 0 | 0 |
| GOCC_INTERCALATED_DISC1 | postnatal | c5_gocc | ExN | -0.3228 | 0.3228 | 0 | 0 |
| GOCC_NUCLEAR_LAMINA1 | postnatal | c5_gocc | ExN | 0.3217 | 0.3217 | 0 | 0 |
| GOCC_L_TYPE_VOLTAGE_GATED_CALCIUM_CI | postnatal | c5_gocc | ExN | -0.3204 | 0.3204 | 0 | 0 |
| GOCC_INTERPHOTORECEPTOR_MATRIX1 | postnatal | c5_gocc | ExN | 0.3198 | 0.3198 | 0 | 0 |
| GOCC_CYTOPLASMIC_SIDE_OF_LYSOSOMAL_I | postnatal | c5_gocc | ExN | 0.3143 | 0.3143 | 0 | 0 |
| GOCC_DISTAL_AXON1 | postnatal | c5_gocc | ExN | -0.3131 | 0.3131 | 0 | 0 |
| GOCC_SPERM_ANNULUS1 | postnatal | c5_gocc | ExN | 0.3092 | 0.3092 | 0 | 0 |
| GOCC_AP_3_ADAPTOR_COMPLEX1 | postnatal | c5_gocc | ExN | 0.3062 | 0.3062 | 0 | 0 |
| GOCC_PHOSPHORYLASE_KINASE_COMPLEX1 | postnatal | c5_gocc | ExN | 0.3058 | 0.3058 | 0 | 0 |
| GOCC_EXTRINSIC_COMPONENT_OF_PLASMA_I | postnatal | c5_gocc | ExN | -0.3053 | 0.3053 | 0 | 0 |
| GOCC_FAR_SIN_STRIPAK_COMPLEX1 | postnatal | c5_gocc | ExN | 0.2964 | 0.2964 | 0 | 0 |
| GOCC_CRD_MEDIATED_MRNA_STABILITY_COM | postnatal | c5_gocc | ExN | 0.2955 | 0.2955 | 0 | 0 |
| GOCC_DNA_RECOMBINASE_MEDIATOR_COMPL | postnatal | c5_gocc | ExN | 0.2928 | 0.2928 | 0 | 0 |
| GOCC_POSTSYNAPTIC_ACTIN_CYTOSKELETON | postnatal | c5_gocc | ExN | 0.2815 | 0.2815 | 0 | 0 |
| GOCC_SCHMIDT_LANTERMAN_INCISURE1 | postnatal | c5_gocc | ExN | 0.2785 | 0.2785 | 0 | 0 |
| GOCC_HOPS_COMPLEX1 | postnatal | c5_gocc | ExN | 0.2782 | 0.2782 | 0 | 0 |
| GOCC_CLATHRIN_COAT_OF_COATED_PIT1 | postnatal | c5_gocc | ExN | 0.2767 | 0.2767 | 0 | 0 |
| GOCC_STEREOCILIUM_BASE1 | postnatal | c5_gocc | ExN | 0.2759 | 0.2759 | 0 | 0 |
| GOCC_NEURON_PROJECTION_TERMINUS1 | postnatal | c5_gocc | ExN | -0.2727 | 0.2727 | 0 | 0 |
| GOCC_I_BAND1 | postnatal | c5_gocc | ExN | -0.2702 | 0.2702 | 0 | 0 |
| GOCC_MLL3_4_COMPLEX1 | postnatal | c5_gocc | ExN | 0.2652 | 0.2652 | 0 | 0 |
| GOCC_ANCHORING_JUNCTION1 | postnatal | c5_gocc | ExN | -0.2649 | 0.2649 | 0 | 0 |
| GOCC_ELONGATOR_HOLOENZYME_COMPLEX1 | postnatal | c5_gocc | ExN | 0.2630 | 0.2630 | 0 | 0 |
| GOCC_CELL_TIP1 | postnatal | c5_gocc | ExN | 0.2600 | 0.2600 | 0 | 0 |
| GOCC_LEADING_EDGE_MEMBRANE1 | postnatal | c5_gocc | ExN | -0.2563 | 0.2563 | 0 | 0 |
| GOCC_F_ACTIN_CAPPING_PROTEIN_COMPLEX | postnatal | c5_gocc | ExN | 0.2546 | 0.2546 | 0 | 0 |
| GOCC_CIS_GOLGI_NETWORK_MEMBRANE1 | postnatal | c5_gocc | ExN | 0.2538 | 0.2538 | 0 | 0 |
| GOCC_MEMBRANE_PROTEIN_COMPLEX1 | postnatal | c5_gocc | ExN | -0.2490 | 0.2490 | 0 | 0 |
| GOCC_NETWORK_FORMING_COLLAGEN_TRIME | postnatal | c5_gocc | ExN | 0.2481 | 0.2481 | 0 | 0 |
| GOCC_EXTRINSIC_COMPONENT_OF_MEMBRAN | postnatal | c5_gocc | ExN | -0.2464 | 0.2464 | 0 | 0 |
| GOCC_PRESYNAPTIC_ACTIVE_ZONE_MEMBRAI | postnatal | c5_gocc | ExN | -0.2445 | 0.2445 | 0 | 0 |
| GOCC_BASEMENT_MEMBRANE_COLLAGEN_TR | postnatal | c5_gocc | ExN | 0.2428 | 0.2428 | 0 | 0 |
| GOCC_SPERM_CONNECTING_PIECE1 | postnatal | c5_gocc | ExN | 0.2415 | 0.2415 | 0 | 0 |
| GOCC_CYTOPLASMIC_REGION1 | postnatal | c5_gocc | ExN | -0.2412 | 0.2412 | 0 | 0 |
| GOCC_PHOSPHATIDYLINOSITOL_3_KINASE_CC | postnatal | c5_gocc | ExN | 0.2337 | 0.2337 | 0 | 0 |
| GOCC_TRAPPII_PROTEIN_COMPLEX1 | postnatal | c5_gocc | ExN | 0.2294 | 0.2294 | 0 | 0 |
| GOCC_NBAF_COMPLEX1 | postnatal | c5_gocc | ExN | 0.2292 | 0.2292 | 0 | 0 |
| GOCC_NUCLEAR_OUTER_MEMBRANE1 | postnatal | c5_gocc | ExN | -0.2221 | 0.2221 | 0 | 0 |
| GOCC_MITOTIC_SPINDLE_MICROTUBULE1 | postnatal | c5_gocc | ExN | 0.2188 | 0.2188 | 0 | 0 |
| GOCC_ATAC_COMPLEX1 | postnatal | c5_gocc | ExN | 0.2177 | 0.2177 | 0 | 0 |
| GOCC_GUANYL_NUCLEOTIDE_EXCHANGE_FAC | postnatal | c5_gocc | ExN | 0.2170 | 0.2170 | 0 | 0 |
| GOCC_FILOPODIUM1 | postnatal | c5_gocc | ExN | -0.2151 | 0.2151 | 0 | 0 |
| GOCC_CYTOPLASMIC_SIDE_OF_ENDOPLASMIC | postnatal | c5_gocc | ExN | 0.2143 | 0.2143 | 0 | 0 |
| GOCC_PERINUCLEAR_ENDOPLASMIC_RETICUL | postnatal | c5_gocc | ExN | 0.2130 | 0.2130 | 0 | 0 |
| GOCC_CUL5_RING_UBIQUITIN_LIGASE_COMPLI | postnatal | c5_gocc | ExN | 0.2115 | 0.2115 | 0 | 0 |
| GOCC_PHOSPHATIDYLINOSITOL_3_KINASE_CC | postnatal | c5_gocc | ExN | 0.2077 | 0.2077 | 0 | 0 |
| GOCC_MICROVESICLE1 | postnatal | c5_gocc | ExN | 0.2071 | 0.2071 | 0 | 0 |
| GOCC_CONTRACTILE_RING1 | postnatal | c5_gocc | ExN | 0.2040 | 0.2040 | 0 | 0 |
| GOCC_CELL_LEADING_EDGE1 | postnatal | c5_gocc | ExN | -0.1970 | 0.1970 | 0 | 0 |
| GOCC_BLEB1 | postnatal | c5_gocc | ExN | 0.1967 | 0.1967 | 0 | 0 |
| GOCC_RNA_POLYMERASE_III_TRANSCRIPTION | postnatal | c5_gocc | ExN | 0.1907 | 0.1907 | 0 | 0 |
| GOCC_PHOSPHATIDYLINOSITOL_3_KINASE_CC | postnatal | c5_gocc | ExN | 0.1896 | 0.1896 | 0 | 0 |
| GOCC_EXOCYTIC_VESICLE1 | postnatal | c5_gocc | ExN | -0.1893 | 0.1893 | 0 | 0 |
| GOCC_KINESIN_COMPLEX1 | postnatal | c5_gocc | ExN | 0.1860 | 0.1860 | 0 | 0 |
| GOCC_MICROFIBRIL1 | postnatal | c5_gocc | ExN | 0.1850 | 0.1850 | 0 | 0 |
| GOCC_FILAMENTOUS_ACTIN1 | postnatal | c5_gocc | ExN | 0.1826 | 0.1826 | 0 | 0 |
| GOCC_H4_HISTONE_ACETYLTRANSFERASE_CC | postnatal | c5_gocc | ExN | 0.1801 | 0.1801 | 0 | 0 |
| GOCC_CUL2_RING_UBIQUITIN_LIGASE_COMPLI | postnatal | c5_gocc | ExN | 0.1778 | 0.1778 | 0 | 0 |
| GOCC_ESC_E_Z_COMPLEX1 | postnatal | c5_gocc | ExN | 0.1776 | 0.1776 | 0 | 0 |
| GOCC_POLYPRENYL_DIPHOSPHATE_SYNTHAS | postnatal | c5_gocc | ExN | 0.1774 | 0.1774 | 0 | 0 |
| GOCC_ENDOPLASMIC_RETICULUM_TUBULAR_N | postnatal | c5_gocc | ExN | 0.1771 | 0.1771 | 0 | 0 |
| GOCC_BASOLATERAL_PLASMA_MEMBRANE1 | postnatal | c5_gocc | ExN | -0.1771 | 0.1771 | 0 | 0 |
| GOCC_AUTOPHAGOSOME_MEMBRANE1 | postnatal | c5_gocc | ExN | 0.1753 | 0.1753 | 0 | 0 |
| GOCC_SYNAPTIC_VESICLE_MEMBRANE1 | postnatal | c5_gocc | ExN | -0.1698 | 0.1698 | 0 | 0 |
| GOCC_CELL_CORTEX1 | postnatal | c5_gocc | ExN | -0.1689 | 0.1689 | 0 | 0 |
| GOCC_H4_H2A_HISTONE_ACETYLTRANSFERAS | postnatal | c5_gocc | ExN | 0.1680 | 0.1680 | 0 | 0 |
| GOCC_M_BAND1 | postnatal | c5_gocc | ExN | -0.1657 | 0.1657 | 0 | 0 |
| GOCC_RPAP3_R2TP_PREFOLDIN_LIKE_COMPL | postnatal | c5_gocc | ExN | 0.1655 | 0.1655 | 0 | 0 |
| GOCC_SWI_SNF_COMPLEX1 | postnatal | c5_gocc | ExN | 0.1650 | 0.1650 | 0 | 0 |
| GOCC_CONTRACTILE_MUSCLE_FIBER1 | postnatal | c5_gocc | ExN | -0.1640 | 0.1640 | 0 | 0 |
| GOCC_CORTICAL_CYTOSKELETON1 | postnatal | c5_gocc | ExN | -0.1596 | 0.1596 | 0 | 0 |
| GOCC_DNA_REPAIR_COMPLEX1 | postnatal | c5_gocc | ExN | 0.1596 | 0.1596 | 0 | 0 |
| GOCC_BASAL_PART_OF_CELL1 | postnatal | c5_gocc | ExN | -0.1576 | 0.1576 | 0 | 0 |
| GOCC_CELL_PROJECTION_MEMBRANE1 | postnatal | c5_gocc | ExN | -0.1569 | 0.1569 | 0 | 0 |
| GOCC_OXOGLUTARATE_DEHYDROGENASE_CC | postnatal | c5_gocc | ExN | 0.1558 | 0.1558 | 0 | 0 |
| GOCC_BBSOME1 | postnatal | c5_gocc | ExN | 0.1555 | 0.1555 | 0 | 0 |
| GOCC_CUL4A_RING_E3_UBIQUITIN_LIGASE_CC | postnatal | c5_gocc | ExN | 0.1539 | 0.1539 | 0 | 0 |
| GOCC_COMPLEX_OF_COLLAGEN_TRIMERS1 | postnatal | c5_gocc | ExN | 0.1497 | 0.1497 | 0 | 0 |
| GOCC_GOLGI_APPARATUS1 | postnatal | c5_gocc | ExN | -0.1467 | 0.1467 | 0 | 0 |
| GOCC_RNA_N6_METHYLADENOSINE_METHYLTI | postnatal | c5_gocc | ExN | 0.1455 | 0.1455 | 0 | 0 |

| | | | | | | | |
|---|---|---|---|---|---|---|---|
| GOCC_SAGA_TYPE_COMPLEX1 | postnatal | c5_gocc | ExN | 0.1360 | 0.1360 | 0 | 0 |
| GOCC_TRANSCRIPTION_ELONGATION_FACTOR | postnatal | c5_gocc | ExN | 0.1352 | 0.1352 | 0 | 0 |
| GOCC_CIS_GOLGI_NETWORK1 | postnatal | c5_gocc | ExN | 0.1343 | 0.1343 | 0 | 0 |
| GOCC_PHAGOPHORE_ASSEMBLY_SITE1 | postnatal | c5_gocc | ExN | 0.1342 | 0.1342 | 0 | 0 |
| GOCC_CELL_SURFACE1 | postnatal | c5_gocc | ExN | -0.1339 | 0.1339 | 0 | 0 |
| GOCC_NUCLEAR_MEMBRANE1 | postnatal | c5_gocc | ExN | -0.1327 | 0.1327 | 0 | 0 |
| GOCC_RECEPTOR_COMPLEX1 | postnatal | c5_gocc | ExN | -0.1321 | 0.1321 | 0 | 0 |
| GOCC_PROTEASOME_COMPLEX1 | postnatal | c5_gocc | ExN | 0.1319 | 0.1319 | 0 | 0 |
| GOCC_CLATHRIN_COATED_PIT1 | postnatal | c5_gocc | ExN | 0.1311 | 0.1311 | 0 | 0 |
| GOCC_LUMENAL_SIDE_OF_ENDOPLASMIC_RET | postnatal | c5_gocc | ExN | 0.1282 | 0.1282 | 0 | 0 |
| GOCC_ACTIN_CYTOSKELETON1 | postnatal | c5_gocc | ExN | -0.1256 | 0.1256 | 0 | 0 |
| GOCC_SMALL_NUCLEAR_RIBONUCLEOPROTEI | postnatal | c5_gocc | ExN | 0.1233 | 0.1233 | 0 | 0 |
| GOCC_CLEAVAGE_FURROW1 | postnatal | c5_gocc | ExN | 0.1229 | 0.1229 | 0 | 0 |
| GOCC_NUCLEAR_ENVELOPE1 | postnatal | c5_gocc | ExN | -0.1222 | 0.1222 | 0 | 0 |
| GOCC_AUTOPHAGOSOME1 | postnatal | c5_gocc | ExN | 0.1219 | 0.1219 | 0 | 0 |
| GOCC_TRANSPORT_VESICLE1 | postnatal | c5_gocc | ExN | -0.1201 | 0.1201 | 0 | 0 |
| GOCC_TRANS_GOLGI_NETWORK1 | postnatal | c5_gocc | ExN | -0.1201 | 0.1201 | 0 | 0 |
| GOCC_ACETYLTRANSFERASE_COMPLEX1 | postnatal | c5_gocc | ExN | 0.1194 | 0.1194 | 0 | 0 |
| GOCC_ACTIN_BASED_CELL_PROJECTION1 | postnatal | c5_gocc | ExN | -0.1181 | 0.1181 | 0 | 0 |
| GOCC_MICROTUBULE_CYTOSKELETON1 | postnatal | c5_gocc | ExN | -0.1161 | 0.1161 | 0 | 0 |
| GOCC_PERINUCLEAR_REGION_OF_CYTOPLASI | postnatal | c5_gocc | ExN | -0.1146 | 0.1146 | 0 | 0 |
| GOCC_GOLGI_ASSOCIATED_VESICLE_MEMBR/ | postnatal | c5_gocc | ExN | 0.1142 | 0.1142 | 0 | 0 |
| GOCC_EXON_EXON_JUNCTION_COMPLEX1 | postnatal | c5_gocc | ExN | 0.1130 | 0.1130 | 0 | 0 |
| GOCC_CATALYTIC_STEP_2_SPLICEOSOME1 | postnatal | c5_gocc | ExN | 0.1081 | 0.1081 | 0 | 0 |
| GOCC_LATE_ENDOSOME_MEMBRANE1 | postnatal | c5_gocc | ExN | 0.1080 | 0.1080 | 0 | 0 |
| GOCC_SUPRAMOLECULAR_COMPLEX1 | postnatal | c5_gocc | ExN | -0.1066 | 0.1066 | 0 | 0 |
| GOCC_GOLGI_APPARATUS_SUBCOMPARTMEN | postnatal | c5_gocc | ExN | -0.1062 | 0.1062 | 0 | 0 |
| GOCC_MITOTIC_SPINDLE_POLE1 | postnatal | c5_gocc | ExN | 0.1061 | 0.1061 | 0 | 0 |
| GOCC_SUPRAMOLECULAR_POLYMER1 | postnatal | c5_gocc | ExN | -0.1037 | 0.1037 | 0 | 0 |
| GOCC_ALPHA_KETOACID_DEHYDROGENASE_( | postnatal | c5_gocc | ExN | 0.1020 | 0.1020 | 0 | 0 |
| GOCC_NUCLEAR_UBIQUITIN_LIGASE_COMPLE) | postnatal | c5_gocc | ExN | 0.1018 | 0.1018 | 0 | 0 |
| GOCC_AP_TYPE_MEMBRANE_COAT_ADAPTOR | postnatal | c5_gocc | ExN | 0.1016 | 0.1016 | 0 | 0 |
| GOCC_CENTROSOME1 | postnatal | c5_gocc | ExN | -0.0984 | 0.0984 | 0 | 0 |
| GOCC_TRANS_GOLGI_NETWORK_TRANSPORT_ | postnatal | c5_gocc | ExN | 0.0974 | 0.0974 | 0 | 0 |
| GOCC_MICROTUBULE_ORGANIZING_CENTER1 | postnatal | c5_gocc | ExN | -0.0966 | 0.0966 | 0 | 0 |
| GOCC_SM_LIKE_PROTEIN_FAMILY_COMPLEX1 | postnatal | c5_gocc | ExN | 0.0953 | 0.0953 | 0 | 0 |
| GOCC_ENDOPEPTIDASE_COMPLEX1 | postnatal | c5_gocc | ExN | 0.0941 | 0.0941 | 0 | 0 |
| GOCC_SWI_SNF_SUPERFAMILY_TYPE_COMPLE | postnatal | c5_gocc | ExN | 0.0939 | 0.0939 | 0 | 0 |
| GOCC_VESICLE_MEMBRANE1 | postnatal | c5_gocc | ExN | -0.0938 | 0.0938 | 0 | 0 |
| GOCC_SPLICEOSOMAL_COMPLEX1 | postnatal | c5_gocc | ExN | 0.0933 | 0.0933 | 0 | 0 |
| GOCC_P_BODY1 | postnatal | c5_gocc | ExN | 0.0917 | 0.0917 | 0 | 0 |
| GOCC_PHOTORECEPTOR_DISC_MEMBRANE1 | postnatal | c5_gocc | ExN | 0.0891 | 0.0891 | 0 | 0 |
| GOCC_CILIARY_MEMBRANE1 | postnatal | c5_gocc | ExN | 0.0887 | 0.0887 | 0 | 0 |
| GOCC_VACUOLAR_LUMEN1 | postnatal | c5_gocc | ExN | 0.0886 | 0.0886 | 0 | 0 |
| GOCC_GOLGI_ASSOCIATED_VESICLE1 | postnatal | c5_gocc | ExN | 0.0881 | 0.0881 | 0 | 0 |
| GOCC_CELL_DIVISION_SITE1 | postnatal | c5_gocc | ExN | 0.0875 | 0.0875 | 0 | 0 |
| GOCC_PEPTIDASE_COMPLEX1 | postnatal | c5_gocc | ExN | 0.0858 | 0.0858 | 0 | 0 |
| GOCC_CONDENSED_NUCLEAR_CHROMOSOME | postnatal | c5_gocc | ExN | 0.0847 | 0.0847 | 0 | 0 |
| GOCC_MICROBODY_LUMEN1 | postnatal | c5_gocc | ExN | 0.0845 | 0.0845 | 0 | 0 |
| GOCC_ENDOSOME1 | postnatal | c5_gocc | ExN | -0.0843 | 0.0843 | 0 | 0 |
| GOCC_BASEMENT_MEMBRANE1 | postnatal | c5_gocc | ExN | 0.0826 | 0.0826 | 0 | 0 |
| GOCC_METHYLTRANSFERASE_COMPLEX1 | postnatal | c5_gocc | ExN | 0.0814 | 0.0814 | 0 | 0 |
| GOCC_SIDE_OF_MEMBRANE1 | postnatal | c5_gocc | ExN | -0.0794 | 0.0794 | 0 | 0 |
| GOCC_APICAL_PART_OF_CELL1 | postnatal | c5_gocc | ExN | -0.0782 | 0.0782 | 0 | 0 |
| GOCC_CUL4_RING_E3_UBIQUITIN_LIGASE_COI | postnatal | c5_gocc | ExN | 0.0782 | 0.0782 | 0 | 0 |
| GOCC_SECRETORY_VESICLE1 | postnatal | c5_gocc | ExN | -0.0760 | 0.0760 | 0 | 0 |
| GOCC_SITE_OF_DOUBLE_STRAND_BREAK1 | postnatal | c5_gocc | ExN | 0.0753 | 0.0753 | 0 | 0 |
| GOCC_PHAGOCYTIC_VESICLE1 | postnatal | c5_gocc | ExN | 0.0732 | 0.0732 | 0 | 0 |
| GOCC_LIPID_DROPLET1 | postnatal | c5_gocc | ExN | 0.0726 | 0.0726 | 0 | 0 |
| GOCC_NUCLEAR_PERIPHERY1 | postnatal | c5_gocc | ExN | 0.0694 | 0.0694 | 0 | 0 |
| GOCC_CENTRIOLAR_SATELLITE1 | postnatal | c5_gocc | ExN | 0.0692 | 0.0692 | 0 | 0 |
| GOCC_SITE_OF_DNA_DAMAGE1 | postnatal | c5_gocc | ExN | 0.0666 | 0.0666 | 0 | 0 |
| GOCC_MITOCHONDRIAL_MATRIX1 | postnatal | c5_gocc | ExN | 0.0665 | 0.0665 | 0 | 0 |
| GOCC_ACTIN_FILAMENT1 | postnatal | c5_gocc | ExN | 0.0660 | 0.0660 | 0 | 0 |
| GOCC_CARDIAC_MYOFIBRIL1 | postnatal | c5_gocc | ExN | 0.0631 | 0.0631 | 0 | 0 |
| GOCC_POLE_PLASM1 | postnatal | c5_gocc | ExN | 0.0621 | 0.0621 | 0 | 0 |
| GOCC_VACUOLAR_MEMBRANE1 | postnatal | c5_gocc | ExN | 0.0584 | 0.0584 | 0 | 0 |
| GOCC_NUCLEAR_BODY1 | postnatal | c5_gocc | ExN | -0.0575 | 0.0575 | 0 | 0 |
| GOCC_MICROVILLUS1 | postnatal | c5_gocc | ExN | 0.0556 | 0.0556 | 0 | 0 |
| GOCC_CHROMOSOME_CENTROMERIC_REGION | postnatal | c5_gocc | ExN | 0.0553 | 0.0553 | 0 | 0 |
| GOCC_COLLAGEN_TRIMER1 | postnatal | c5_gocc | ExN | 0.0550 | 0.0550 | 0 | 0 |
| GOCC_POLYMERIC_CYTOSKELETAL_FIBER1 | postnatal | c5_gocc | ExN | -0.0536 | 0.0536 | 0 | 0 |
| GOCC_ENDOPLASMIC_RETICULUM_LUMEN1 | postnatal | c5_gocc | ExN | 0.0528 | 0.0528 | 0 | 0 |
| GOCC_ORGANELLE_ENVELOPE1 | postnatal | c5_gocc | ExN | -0.0509 | 0.0509 | 0 | 0 |
| GOCC_NUCLEAR_CHROMOSOME1 | postnatal | c5_gocc | ExN | 0.0506 | 0.0506 | 0 | 0 |
| GOCC_ENDOPLASMIC_RETICULUM_PROTEIN_C | postnatal | c5_gocc | ExN | 0.0498 | 0.0498 | 0 | 0 |
| GOCC_NUCLEAR_PORE1 | postnatal | c5_gocc | ExN | 0.0494 | 0.0494 | 0 | 0 |
| GOCC_CONDENSED_CHROMOSOME_CENTROM | postnatal | c5_gocc | ExN | 0.0493 | 0.0493 | 0 | 0 |
| GOCC_GOLGI_MEMBRANE1 | postnatal | c5_gocc | ExN | -0.0466 | 0.0466 | 0 | 0 |
| GOCC_CONDENSED_CHROMOSOME1 | postnatal | c5_gocc | ExN | 0.0452 | 0.0452 | 0 | 0 |
| GOCC_ACROSOMAL_VESICLE1 | postnatal | c5_gocc | ExN | 0.0442 | 0.0442 | 0 | 0 |
| GOCC_COLLAGEN_CONTAINING_EXTRACELLUI | postnatal | c5_gocc | ExN | 0.0430 | 0.0430 | 0 | 0 |
| GOCC_PROTEIN_DNA_COMPLEX1 | postnatal | c5_gocc | ExN | -0.0428 | 0.0428 | 0 | 0 |
| GOCC_BLOOD_MICROPARTICLE1 | postnatal | c5_gocc | ExN | 0.0418 | 0.0418 | 0 | 0 |
| GOCC_MEMBRANE_MICRODOMAIN1 | postnatal | c5_gocc | ExN | -0.0402 | 0.0402 | 0 | 0 |
| GOCC_RNA_POLYMERASE_COMPLEX1 | postnatal | c5_gocc | ExN | 0.0348 | 0.0348 | 0 | 0 |
| GOCC_CHROMOSOMAL_REGION1 | postnatal | c5_gocc | ExN | 0.0345 | 0.0345 | 0 | 0 |
| REACTOME_FORMATION_OF_APOPTOSOME1 | postnatal | keywords_c2 | Non-neuron | 0.0572 | 0.0572 | 0.0003 | 0.0450 |
| GRAESSMANN_APOPTOSIS_BY_DOXORUBICIN_ | postnatal | keywords_c2 | Non-neuron | 0.0088 | 0.0088 | 0.0001 | 0.0150 |

| | | | | | | | | |
|---|---|---|---|---|---|---|---|---|
| KEGG_MEDICUS_VARIANT_MUTATION_CAUSED | postnatal | keywords_c2 | ExN | -0.6393 | | 0.6393 | 0.0001 | 0.0150 |
| KEGG_MEDICUS_REFERENCE_MGLUR5_CA2_A | postnatal | keywords_c2 | ExN | -0.6175 | | 0.6175 | 0.0002 | 0.0300 |
| KEGG_MEDICUS_VARIANT_MUTATION_CAUSED | postnatal | keywords_c2 | ExN | -0.5462 | | 0.5462 | 0 | 0 |
| KEGG_MEDICUS_REFERENCE_AUTOPHAGOSOI | postnatal | keywords_c2 | ExN | 0.4153 | | 0.4153 | 0 | 0 |
| REACTOME_SUPPRESSION_OF_APOPTOSIS1 | postnatal | keywords_c2 | ExN | 0.3188 | | 0.3188 | 0.0001 | 0.0150 |
| REACTOME_SARS_COV_2_MODULATES_AUTOI | postnatal | keywords_c2 | ExN | 0.3043 | | 0.3043 | 0 | 0 |
| KEGG_MEDICUS_REFERENCE_AUTOPHAGY_VE | postnatal | keywords_c2 | ExN | 0.3002 | | 0.3002 | 0.0001 | 0.0150 |
| KEGG_MEDICUS_REFERENCE_AUTOPHAGY_VE | postnatal | keywords_c2 | ExN | 0.2987 | | 0.2987 | 0 | 0 |
| RAMJAUN_APOPTOSIS_BY_TGFB1_VIA_SMAD4 | postnatal | keywords_c2 | ExN | 0.2978 | | 0.2978 | 0 | 0 |
| REACTOME_TP53_REGULATES_TRANSCRIPTIO | postnatal | keywords_c2 | ExN | 0.2908 | | 0.2908 | 0 | 0 |
| REACTOME_TP53_REGULATES_TRANSCRIPTIO | postnatal | keywords_c2 | ExN | 0.2442 | | 0.2442 | 0 | 0 |
| REACTOME_REGULATION_OF_MITF_M_DEPENI | postnatal | keywords_c2 | ExN | 0.2436 | | 0.2436 | 0.0001 | 0.0150 |
| REACTOME_CASPASE_MEDIATED_CLEAVAGE_ | postnatal | keywords_c2 | ExN | 0.2383 | | 0.2383 | 0 | 0 |
| REACTOME_FORMATION_OF_APOPTOSOME1 | postnatal | keywords_c2 | ExN | 0.2358 | | 0.2358 | 0 | 0 |
| REACTOME_ONCOGENE_INDUCED_SENESCEN( | postnatal | keywords_c2 | ExN | 0.2326 | | 0.2326 | 0.0001 | 0.0150 |
| GALI_TP53_TARGETS_APOPTOTIC_UP1 | postnatal | keywords_c2 | ExN | 0.2237 | | 0.2237 | 0 | 0 |
| KEGG_MEDICUS_REFERENCE_C9ORF72_MEDIA | postnatal | keywords_c2 | ExN | 0.2180 | | 0.2180 | 0.0003 | 0.0450 |
| CHICAS_RB1_TARGETS_SENESCENT1 | postnatal | keywords_c2 | ExN | -0.2057 | | 0.2057 | 0 | 0 |
| KEGG_MEDICUS_REFERENCE_AUTOPHAGY_VE | postnatal | keywords_c2 | ExN | 0.1905 | | 0.1905 | 0 | 0 |
| WP_APOPTOSIS_MODULATION_BY_HSP701 | postnatal | keywords_c2 | ExN | -0.1779 | | 0.1779 | 0.0001 | 0.0150 |
| WP_AUTOPHAGY_IN_PANCREATIC_DUCTAL_AD | postnatal | keywords_c2 | ExN | 0.1735 | | 0.1735 | 0.0003 | 0.0450 |
| KEGG_MEDICUS_REFERENCE_AUTOPHAGY_VE | postnatal | keywords_c2 | ExN | 0.1700 | | 0.1700 | 0.0001 | 0.0150 |
| WP_PERTURBATIONS_TO_HOSTCELL_AUTOPH | postnatal | keywords_c2 | ExN | 0.1612 | | 0.1612 | 0 | 0 |
| WP_HOSTPATHOGEN_INTERACTION_OF_HUMA | postnatal | keywords_c2 | ExN | 0.1470 | | 0.1470 | 0.0001 | 0.0150 |
| KEGG_MEDICUS_VARIANT_MUTATION_CAUSED | postnatal | keywords_c2 | ExN | 0.1436 | | 0.1436 | 0.0003 | 0.0450 |
| KEGG_MEDICUS_REFERENCE_AUTOPHAGY_VE | postnatal | keywords_c2 | ExN | 0.1418 | | 0.1418 | 0.0003 | 0.0450 |
| KEGG_REGULATION_OF_AUTOPHAGY1 | postnatal | keywords_c2 | ExN | 0.1416 | | 0.1416 | 0 | 0 |
| HAMAI_APOPTOSIS_VIA_TRAIL_UP1 | postnatal | keywords_c2 | ExN | -0.1282 | | 0.1282 | 0 | 0 |
| REACTOME_DNA_DAMAGE_TELOMERE_STRES | postnatal | keywords_c2 | ExN | 0.1236 | | 0.1236 | 0 | 0 |
| WU_APOPTOSIS_BY_CDKN1A_NOT_VIA_TP531 | postnatal | keywords_c2 | ExN | 0.1217 | | 0.1217 | 0 | 0 |
| WP_AUTOPHAGY1 | postnatal | keywords_c2 | ExN | 0.1196 | | 0.1196 | 0.0001 | 0.0150 |
| GRAESSMANN_APOPTOSIS_BY_DOXORUBICIN_ | postnatal | keywords_c2 | ExN | -0.1189 | | 0.1189 | 0 | 0 |
| WP_NANOPARTICLE_TRIGGERED_AUTOPHAGI( | postnatal | keywords_c2 | ExN | 0.1105 | | 0.1105 | 0.0001 | 0.0150 |
| REACTOME_APOPTOTIC_CLEAVAGE_OF_CELL | postnatal | keywords_c2 | ExN | 0.1101 | | 0.1101 | 0 | 0 |
| KEGG_MEDICUS_REFERENCE_AUTOPHAGOSOI | postnatal | keywords_c2 | ExN | 0.1055 | | 0.1055 | 0.0001 | 0.0150 |
| REACTOME_SENESCENCE_ASSOCIATED_SECF | postnatal | keywords_c2 | ExN | 0.0978 | | 0.0978 | 0 | 0 |
| WP_SENESCENCEASSOCIATED_SECRETORY_F | postnatal | keywords_c2 | ExN | 0.0978 | | 0.0978 | 0.0001 | 0.0150 |
| REACTOME_APOPTOTIC_EXECUTION_PHASE1 | postnatal | keywords_c2 | ExN | 0.0965 | | 0.0965 | 0.0002 | 0.0300 |
| WP_CLOCKCONTROLLED_AUTOPHAGY_IN_BON | postnatal | keywords_c2 | ExN | 0.0963 | | 0.0963 | 0 | 0 |
| WU_APOPTOSIS_BY_CDKN1A_VIA_TP531 | postnatal | keywords_c2 | ExN | 0.0889 | | 0.0889 | 0.0001 | 0.0150 |
| REACTOME_LATE_ENDOSOMAL_MICROAUTOPI | postnatal | keywords_c2 | ExN | 0.0853 | | 0.0853 | 0.0003 | 0.0450 |
| WP_TNFRELATED_WEAK_INDUCER_OF_APOPT | postnatal | keywords_c2 | ExN | 0.0791 | | 0.0791 | 0.0001 | 0.0150 |
| WP_APOPTOSIS_MODULATION_AND_SIGNALIN | postnatal | keywords_c2 | ExN | 0.0769 | | 0.0769 | 0 | 0 |
| WP_OMEGA6FATTY_ACIDS_IN_SENESCENCE1 | postnatal | keywords_c2 | ExN | 0.0745 | | 0.0745 | 0 | 0 |
| CONCANNON_APOPTOSIS_BY_EPOXOMICIN_DI | postnatal | keywords_c2 | ExN | -0.0690 | | 0.0690 | 0.0001 | 0.0150 |
| GRAESSMANN_APOPTOSIS_BY_DOXORUBICIN_ | postnatal | keywords_c2 | ExN | 0.0494 | | 0.0494 | 0 | 0 |
| REACTOME_CELLULAR_SENESCENCE1 | postnatal | keywords_c2 | ExN | 0.0473 | | 0.0473 | 0.0001 | 0.0150 |
